# A catalogue of omics biological ageing clocks reveals substantial commonality and associations with disease risk

**DOI:** 10.1101/2021.02.01.429117

**Authors:** Erin Macdonald-Dunlop, Nele Taba, Lucija Klaric, Azra Frkatovic, Rosie Walker, Caroline Hayward, Tonu Esko, Chris Haley, Krista Fischer, James F Wilson, Peter K Joshi

**Author notes:** These Authors contributed equally (James F Wilson, Peter K Joshi).

## Abstract

Biological age (BA), a measure of functional capacity and prognostic of health outcomes that discriminates between individuals of the same chronological age (chronAge), has been estimated using a variety of biomarkers. Previous comparative studies have mainly used epigenetic models (clocks), we use ~1000 participants to create eleven omics ageing clocks, with correlations of 0.45-0.97 with chronAge, even with substantial sub-setting of biomarkers. These clocks track common aspects of ageing with 94% of the variance in chronAge being shared among clocks. The difference between BA and chronAge - omics clock age acceleration (OCAA) - often associates with health measures. One year’s OCAA typically has the same effect on risk factors/10-year disease incidence as 0.46/0.45 years of chronAge. Epigenetic and IgG glycomics clocks appeared to track generalised ageing while others capture specific risks. We conclude BA is measurable and prognostic and that future work should prioritise health outcomes over chronAge.

## Introduction

Age is a phenotype that we are all familiar with, and is a major risk factor for numerous diseases including the largest causes of mortality^1^. We all become acquainted with visible changes that accompany ageing, such as greying hair, baldness, loss of skin elasticity and worsening of posture, and that these vary noticeably amongst individuals of the same chronological age (chronAge). However, there are also molecular hallmarks of ageing such as telomere shortening, genomic instability and cellular senescence that also show variation in individuals of the same chronAge^1^. It has previously been hypothesised that an underlying biological age (BA), likely tagged by these molecular hallmarks, is what gives rise to age-related disease risk^2^. Measuring BA therefore has the potential to be more prognostic of health and functional capacity than chronAge and, as importantly, BA may be reversible^3^, unlike chronAge^4^.

Since this concept was proposed, there has been a push to construct models of BA, using a variety of both statistical methods and types of biomarkers; the resultant estimates we shall term omics clock ages (OCAs). The first OCAs were epigenetic clocks that used methylation levels of CpG sites across the genome - DNA methylation (DNAme) - to estimate chronAge using penalised regression^5,6^. The excess of OCA over chronAge being omics clock age acceleration (OCAA), hopefully measuring an underlying biological effect. DNAme’s verification as a meaningful BA measure, rather than a mere statistical artefact, was confirmed when DNAme OCAA as calculated by Horvath’s clock was shown to be associated with all-cause mortality^7^. Ageing clocks trained on chronAge have since been constructed using DNA methylation^5,6,8,9^, telomere length^9,10^, facial morphology^11^, neuroimaging data^12–15^, metabolomics^9,16–18^, glycomics^19^, proteomics^9,20–22^ and immune cell counts^23^. There has however, been limited comparison of the performance, for example accuracy and correlation, of different omics ageing clocks, particularly in the same set of individuals.

Moreover, there has been inadequate additional progress in demonstrating that the various OCA measures are actually tracking underlying BA beyond chronAge, and whether some clocks’ OCAAs are more aligned to certain outcomes than others. For example, few significant associations of chronAge-trained OCAAs have been found with health outcomes other than mortality and those that do have low effect sizes^24–27^.

The deep omic and health outcome annotation of the Scottish population-based Orkney Complex Disease Study^28^ cohort (ORCADES) permits interrogation of the utility and limitations of BA clocks. Here, we compare the performance of 11 ageing clocks built from 9 different omics assays in the same set of approximately 1000 individuals in ORCADES, including whole body imaging and a clock based on the grand union of all the omics. Next, we assess the biological meaningfulness of the derived OCAA measures, by assessing their association with health-related phenotypes and incident hospital admissions (post-assessment) over up to 10 years follow-up.

The notion of BA raises fundamental questions. Is there one BA for a person, or a set of BAs, perhaps relating to different bodily systems^22,29^. Are measured (chronAge trained) OCAs tracking a single BA, with differences arising due to their focus and accuracy, or are they tracking different underlying BAs? This study aims to shed some light on these issues.

## Results

### Performance of Omics Clocks

We constructed eleven ageing clocks, training on chronAge, in the ORCADES cohort from assays already understood to be able to form effective ageing clocks^5,6,19,20^, covering plasma Immunoglobulin G (IgG) glycans, proteins, metabolites, lipids, DNA methylation and a collection of commonly used clinical measures (such as weight, blood pressure, fasting glucose, etc), which we label Clinomics. To this we added two novel omics sets for clock construction: a DEXA whole body imaging set of body composition measures, and one based on all the omics assays considered simultaneously, which we term Mega-omics, as listed in Table 1 (see Methods for assay descriptions). Rather than creating completely novel DNAme clocks when effective and extensively studied published clocks exist, our methylation clocks’ potential predictor sets are the subsets of the CpG sites used in Hannum and Horvath’s epigenetic clocks available on the Illumina EPIC 850k methylation array. With this caveat, all clocks were derived from scratch using the set of available predictors and elastic net regression.

**Table 1.** Multiple omics make accurate ageing clocks. Indicating for each omics assay: N Individuals: the number of individuals in the ORCADES cohort that passes quality control, N Predictors Available: the number of predictors passing assay-level quality control and therefore available for selection for inclusion in the standard model, N Predictors Selected: the number of predictors selected for inclusion in the standard model, r: Pearson correlation of omics clock age (OCA) and chronAge. DEXA: Dual X-ray absorptiometry, DNAme: DNA methylation, CpG: cytosine nucleotide followed by guanine (5’ to 3’ direction), MS: mass spectrometry, NMR: nuclear magnetic resonance, PEA: proximity extension assay, UPLC: ultra-performance liquid chromatography, IgG: Immunoglobulin G. Within each omic category, subject mean age at baseline was 53-56 (SD~15) with an age range across clocks of 16-100, whilst the proportion female ranged from 55-61% (Supplementary Table 1).

| Omic | N Individuals | N Predictors Available | N Predictors Selected | r |
| --- | --- | --- | --- | --- |
| MS Fatty Acids Lipidomics | 952 | 33 | 27 | 0.45 |
| DEXA | 1158 | 28 | 28 | 0.66 |
| MS Complex Lipidomics | 940 | 908 | 130 | 0.7 |
| NMR Metabolomics | 1643 | 86 | 81 | 0.74 |
| UPLC IgG Glycomics | 1937 | 77 | 50 | 0.74 |
| Clinomics | 1815 | 13 | 12 | 0.8 |
| MS Metabolomics | 861 | 682 | 181 | 0.81 |
| DNAme Horvath CpGs | 957 | 333 | 155 | 0.93 |
| PEA Proteomics | 805 | 886 | 203 | 0.93 |
| DNAme Hannum CpGs | 1033 | 62 | 50 | 0.96 |
| Mega Omics | 796 | 2471 | 214 | 0.97 |

We first assessed various forms of penalised regression: LASSO, elastic net with a fixed alpha of 0.5 and elastic net with alpha calculated via cross-validation, training clocks in 75% of the ORCADES cohort and evaluating in the remaining 25% (the testing sample). We found that clock performance in estimating chronAge was independent of penalised regression method used, across all the assays (Supplementary Figure 1) and so elastic net regression with a fixed alpha of 0.5 only was employed in subsequent analyses.

Ages estimated by the model in the test set (i.e. OCAs) were highly correlated with chronAge for the majority of the omics clocks tested (Table 1), particularly PEA proteomics (r=0.93) and DNAme based (r=0.96 Hannum CpGs, r=0.93 Horvath CpGs) clocks (correlations in the training set in Supplementary Figure 2). Unsurprisingly, the mega-omics OCA had the highest correlation (r=0.97). Although all features were given equal opportunity to contribute to the mega-omics clock, those selected by the regression were predominantly DNAme- and PEA proteomics-based (34.6% CpGs, 31.8% PEA Proteomics, 20.6% MS metabolites, 13.1% other). We found that the MS Fatty Acids Lipidomics OCA had the lowest correlation with chronAge (r=0.45; Figure 1). The number of biomarkers available and then selected for model inclusion for each omics clock are indicated in Table 1 (Full list of biomarkers measured in each assay in Supplementary Table 2 and coefficients for all clocks in Supplementary Table 3).

**Figure 1.**
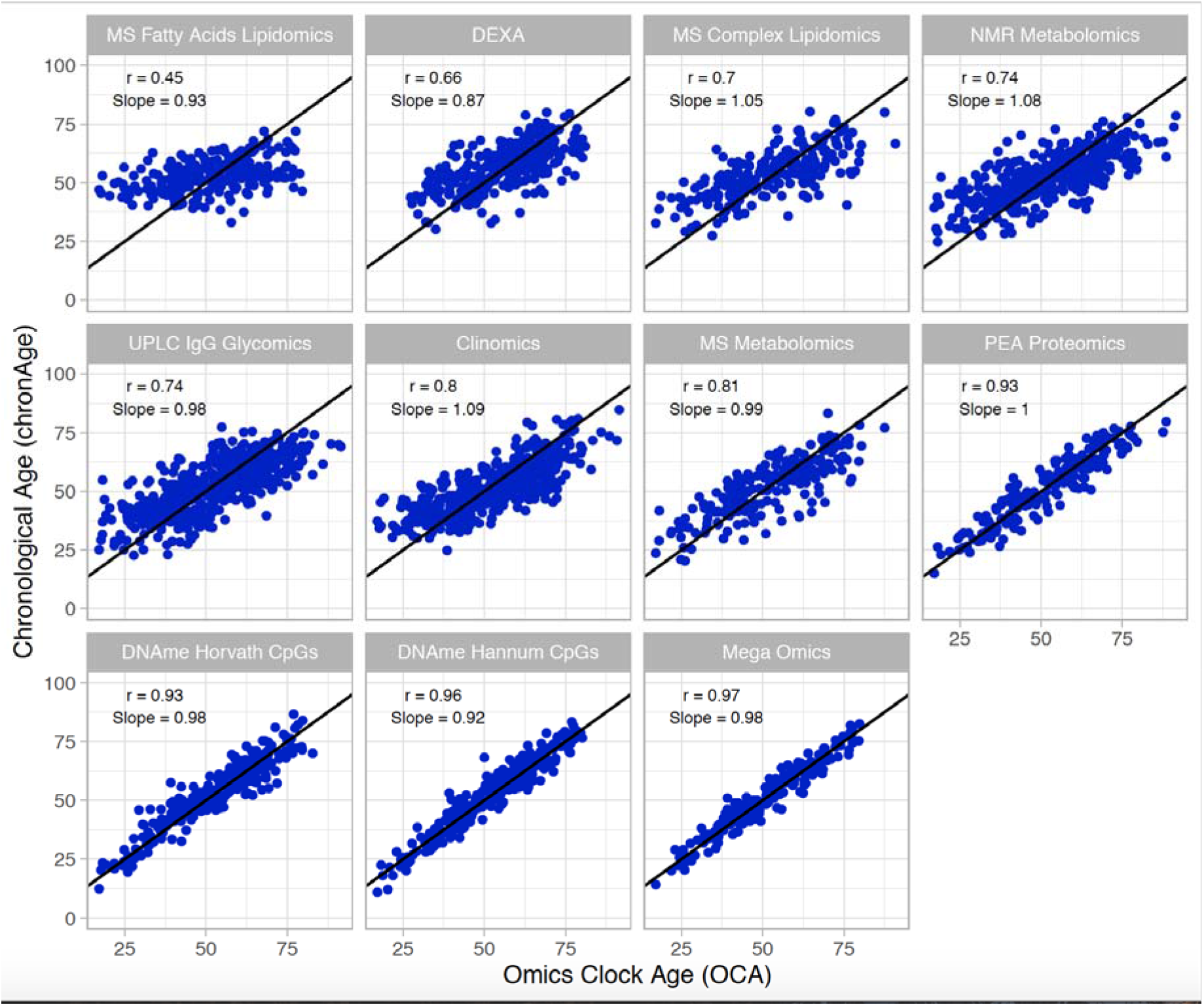
Multiple omics estimate chronological age, to varying degrees of accuracy, in a broadly unbiased manner. The correlations of chronAge on the y-axis with ages estimated by the omics ageing clock (OCA) in the ORCADES testing sample. Pearson correlation coefficient (r) and the slope of the regression of OCA on chronAge are indicated in each panel. Identity line indicated in black.

### Validation of Clock Performance in Independent Cohorts

We next used the clocks trained in ORCADES to estimate age in independent European cohorts to validate if they were more widely applicable beyond the Orkney population. We found that correlations between OCA and chronAge replicated to varying degrees in independent populations (Supplementary Figure 3). PEA proteomics and DNAme based clocks produced correlations of OCA and chronAge in the range of 0.89-0.98 in European cohorts replicating the range of 0.91-0.96 in ORCADES. UPLC IgG glycomics and Clinomics OCAs in independent populations showed a range of OCA-chronAge correlations of 0.56-0.62 compared to the 0.74-0.80 in ORCADES. Whilst the NMR metabolomics and DEXA did not replicate with correlations of 0.26-0.55 in validation cohorts compared with 0.66-0.73 in ORCADES.

### Accurate Performance of Clocks with Substantial Core Subset of Biomarkers

If the aim is to create BA clocks that have the potential to be clinically useful, it would be more efficient and cost effective to reduce the numbers of biomarkers that need to be measured in patients. To this end, we investigated the performance of our clocks using a reduced set of biomarkers. For each of our 11 omics clocks a “core” clock was constructed using only those biomarkers which were selected for model inclusion in >95% of 500 iterations of our clock construction procedure, as done by Enroth *et al*.^20^ (See Methods for details). Comparable correlations of OCA and chronAge were achieved across all 11 clocks with a substantial subset or core of biomarkers (Figure 2), highlighting the potential for accurate OCAs with a small number of predictors (e.g. 30s-60s of biomarkers).

**Figure 2.**
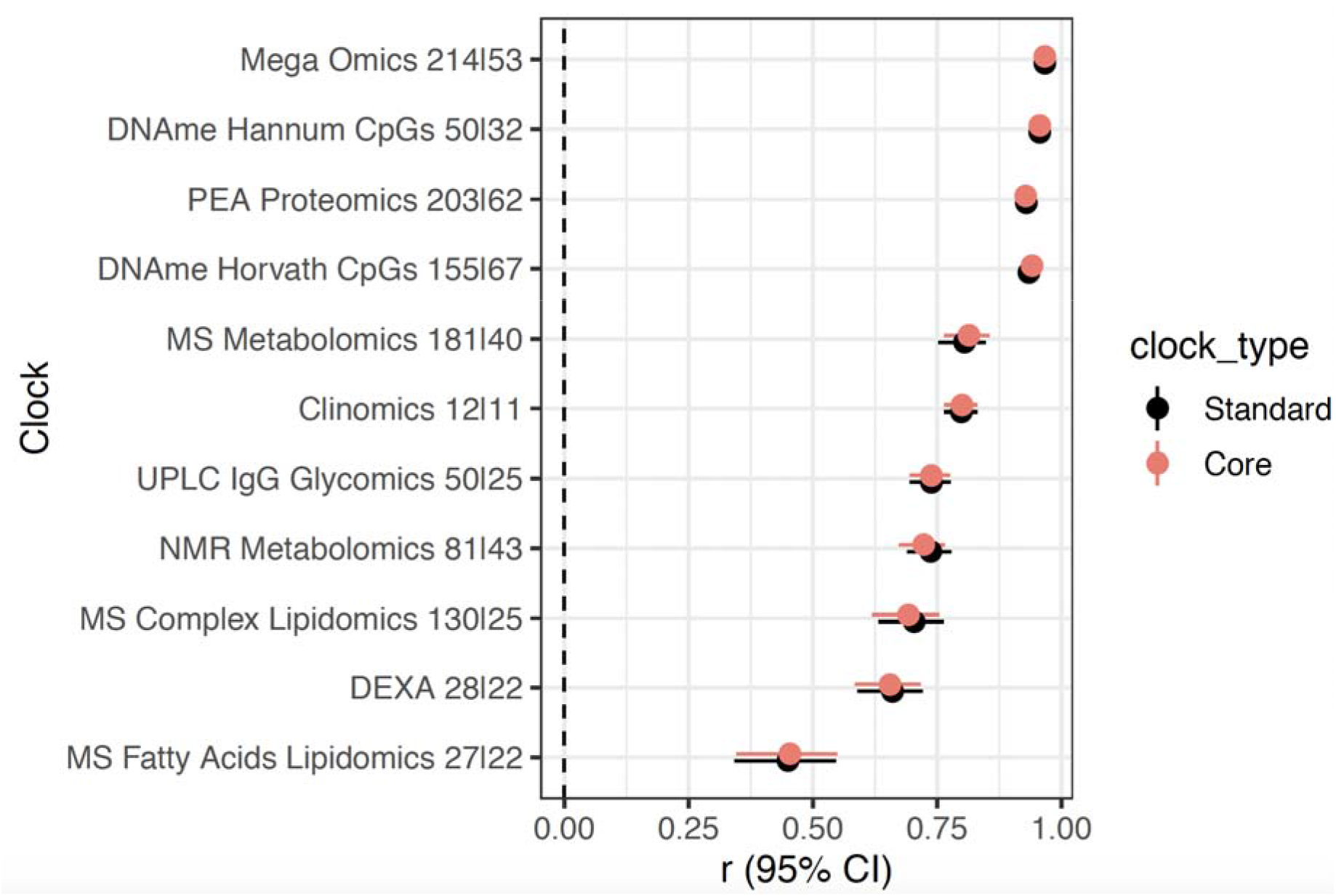
Substantial subsetting of biomarkers results in little dilution of accuracy. Pearson’s correlation (r) and 95% confidence interval of chronAge and OCAs from standard and core models for each omics assay indicated on the y-axis in the ORCADES testing sample. The number of predictors selected for inclusion in the standard and then core models are indicates in the y-axis labels (standardļcore).

### Comparison of Biological Age Between Clocks

Omics Clock Age Accelerations (OCAAs) showed varying degrees of positive correlation between clocks (Figure 3). Unsurprisingly, the two DNAme based OCAAs were the most correlated with each other (r=0.73) and, in hierarchical clustering, formed a group on their own. The four clocks that are primarily constructed from lipid species and fractions, MS Fatty Acids Lipidomics, MS Complex lipidomics, NMR Metabolomics and MS Metabolomics clocks, all clustered together. The DEXA, Clinomics and UPLC IgG glycomics clocks formed a related group. Interestingly, the PEA Proteomics OCAA clustered between the DNAme and glycomics-DEXA-Clinomics-lipidomics cluster, on its own.

**Figure 3.**
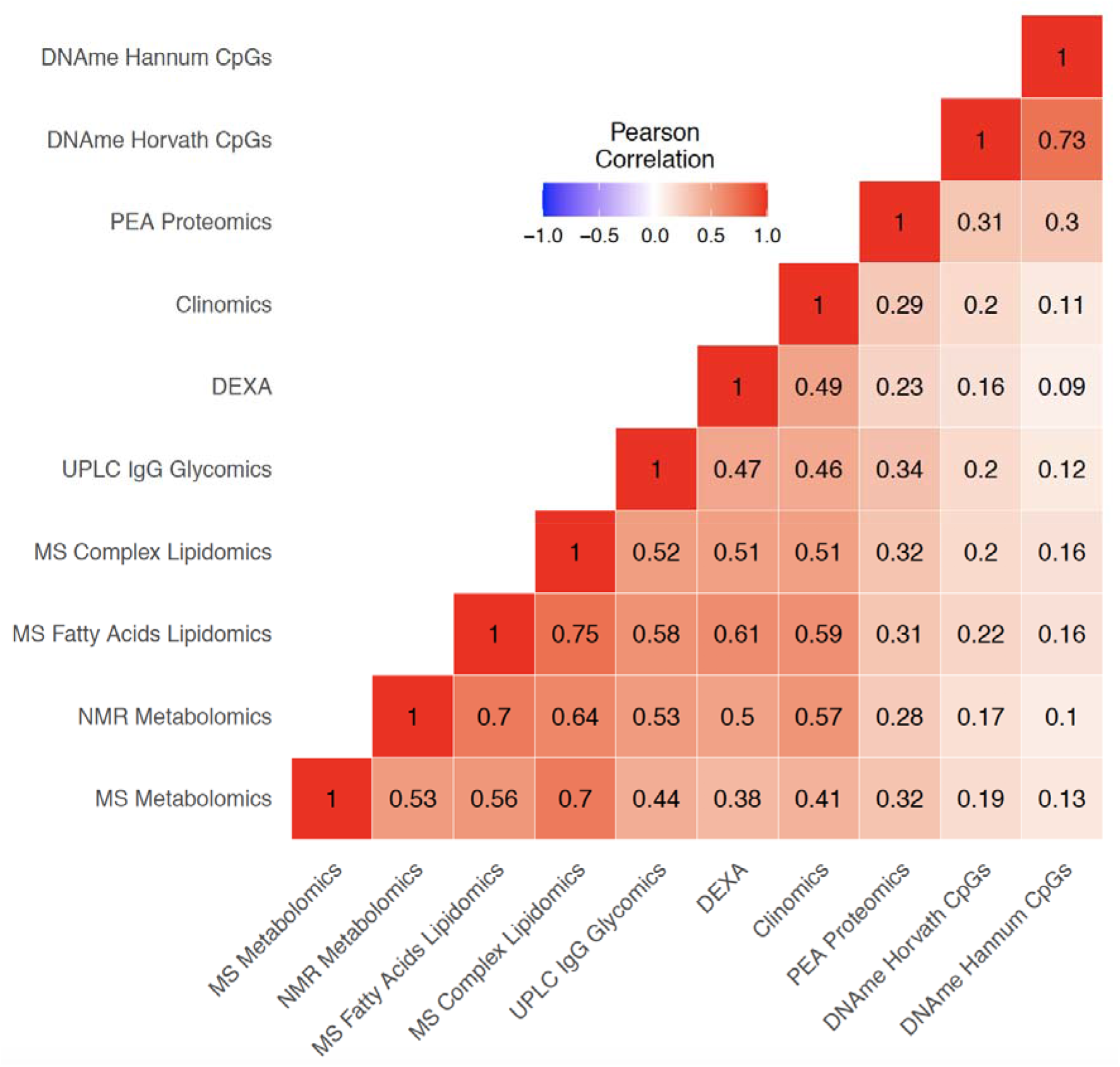
Variable positive correlations between different omics age accelerations. Pearson correlation of OCAAs (omics clock age-chronAge) in ORCADES testing and training samples. Colour indicates the direction and the shade and number indicates the magnitude of the correlation. Rows and columns are ordered based on hierarchical clustering of the pairwise correlations.

### Proportions of Variance in Age Explained by Different Clocks

To determine if our different clocks are explaining the same or different variance in chronAge, we partitioned the variance in chronAge explained among our clocks. We calculated the unique variance in chronAge explained by each OCA as the squared part correlations of chronAge and OCA, while controlling for all other clocks. 93.9% of the variance in chronAge is explained by two or more clocks whilst 4.1% remains unexplained by the 10 ageing clocks tested, with the remaining 1.9% being explained by one clock uniquely (Supplementary Figure 4a). The PEA proteomics and Hannum CpG clocks explain the most variance in chronAge uncaptured by any other clock (0.59% and 0.46% respectively; Supplementary Figure 4b). Pairwise clock comparisons are shown in Supplementary Figure 5.

Having found that clocks overlap in the information they provide about chronAge, we tested to see if, together, pairs of clocks jointly explained a different proportion of variance in chronAge than would be expected if the clocks were each independently sampling from a latent set of complete predictors of chronAge (ISLSP). This analysis should reveal whether the clocks were tracking complementary dimensions of ageing: situations where the pair of clocks overlapped less than expected if they were independently sampling (negative values on this scale). Strikingly, excess overlap was found across all pairs of clocks (Figure 4), with the lowest excess overlap value measured at 0.41 (comparison of the NMR Metabolomics and DEXA clocks): all 10 omics clocks, considered pairwise, track more common rather than complementary aspects of chronAge.

**Figure 4.**
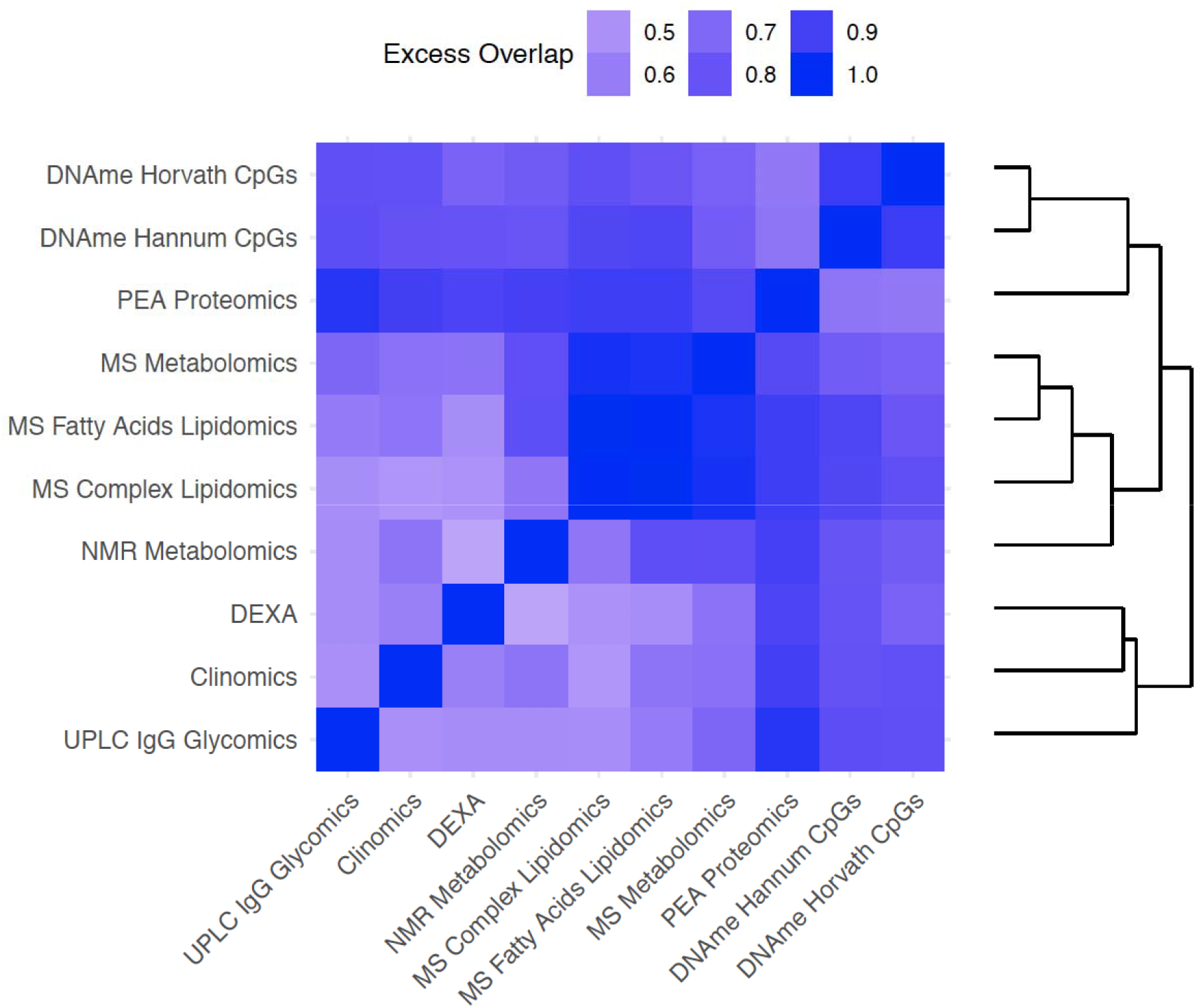
Bivariate analyses reveal that clock pairs tend to overlap more than expected by chance in the variance in ChronAge they explain. The amount of excess overlap than would be expected by chance is indicated for each pair of clocks. This is the deviation of the observed variance in chronAge explained by a bivariate model containing a pair of OCAs and the variance expected to be explained by that pair given that we know how much variance in chronAge they explain individually, if each of the clocks were independent samples from a set of latent complete predictors. This measure of deviation of observed from expected is scaled (See Methods for details) so that a value of 1 means that the second clock is adding no more information than the first, meaning that they overlap entirely in the information they provide about chronAge. A value of 0 would indicate the observed variance explained in chronAge is exactly what is expected if the two clocks were independently sampling. Negative values are possible on this scale but are not observed and would indicate disproportionately complementary components of chronAge were being tracked.

The most overlapping were the MS Fatty Acids Lipidomics and the MS Complex Lipidomics clocks (excess overlap of 0.98; note on our scale, a clock shows 1.00 excess overlap with itself, whilst ISLSP would show 0.00). These two clocks formed a cluster with the clocks derived from NMR and MS Metabolomics (which both contain many lipid features). Similarly, the two DNAme-based clocks clustered tightly together with an excess overlap of 0.91. As these clocks are extremely accurate, a large amount of overlap in variance explained is inevitable; they are tracking common aspects of ageing.

### OCAAs compared to chronAge as predictors of disease risk

We next sought to test the effect of OCAAs compared to chronAge on risk factors and post assessment disease incidence, as measured by hospitalisation in the ORCADES cohort, where the outcome was thought *a priori* to associate with age. For risk factors we chose body mass index (BMI), systolic blood pressure (SBP), cortisol, creatinine, C-reactive protein (CRP), forced expiratory volume in 1 second (FEV1), Framingham Risk Score, and total cholesterol. For diseases we chose five International Statistical Classification of Diseases and Related Health Problems(ICD)-10 Chapters: II (Neoplasms - codes C), IV (Endocrine, nutritional and metabolic diseases - codes E), IX (Diseases of the circulatory system - codes I), and X (Diseases of the respiratory system - codes J). The ICD-10 blocks used and their codings are listed in Supplementary Table 4).

In order to compare OCAA and chronAge, we first quantified the effect of chronAge on risk factors and disease (Supplementary Figures 6a & 6b). All 8 risk factors and 32/44 disease blocks were taken forward as they were significantly associated with chronAge (beta>0, FDR<10%) and had >5 incident cases (disease blocks). The effect of chronAge on (standardised) risk factors appeared to vary by trait, whereas for diseases, it appeared that the effect of chronAge (on the hazard ratio scale) might be similar across diseases, with a consistent doubling of risk every 14 years.

We tested for risk factor and disease associations with OCAA, using chronAge and sex as covariates. Results were then rescaled to be per year of chronAge effect by dividing the observed effect of OCAA by the effect of chronAge on the outcomes as identified at the previous step. This was taken trait-by-trait for risk factors, and a single effect for all disease groups and chapters: −0.0492 log_e_ HR.

Despite limited power for detecting OCAA-disease associations, 6/352 tests were statistically significant (FDR<10%) as were 31/88 OCAA-risk factor associations. We also found evidence of enrichment of positive effects of OCAA on both risk factors (85%) and disease (74%), with 35% and 23% being nominally significant (one sided p<0.05), respectively. Across clocks, the inverse variance-weighted mean effect of one year of OCAA on risk factors/disease was the same as 0.45/0.46 years of chronAge (SE~0.01, note here and elsewhere ~ denotes indicative, see Methods for details). For risk factors, as might be expected, this was strongly influenced by an average effect of 1.23 years for Clinomics OCAA (0.16 without Clinomics). Interestingly, the mean effect across all diseases of one year’s DNAme Hannum/Horvath CpGs OCAA was similar to one year of chronAge (ratio: 1.03/0.85, SEs ~0.18), but the effect on risk factors was much lower (ratio: −0.03/-0.01, SEs ~ 0.06). Complete results are shown in Supplementary Table 5 and inverse variance-weighted effects are shown in Supplementary Figure 7b.

In general, only associations with the Clinomics OCAA passed FDR, however both DNAme OCAAs and the UPLC IgG Glycomics OCAA were nominally associated with eleven ICD10 blocks, one more than Clinomics (Figure 5). In contrast, the PEA proteomics clock (r=0.93 with chronAge) showed only one nominally significant disease:OCAA association. Looking at disease groupings, E70-E90 Metabolic disorders and J09-J18 Influenza and Pneumonia showed the most nominal associations across all OCAAs. Curiously, on the other hand, C34-C44 Melanoma and C51-59 Malignant Neoplasms of the female genital organs, showed generally negative associations with OCAAs.

**Figure 5.**
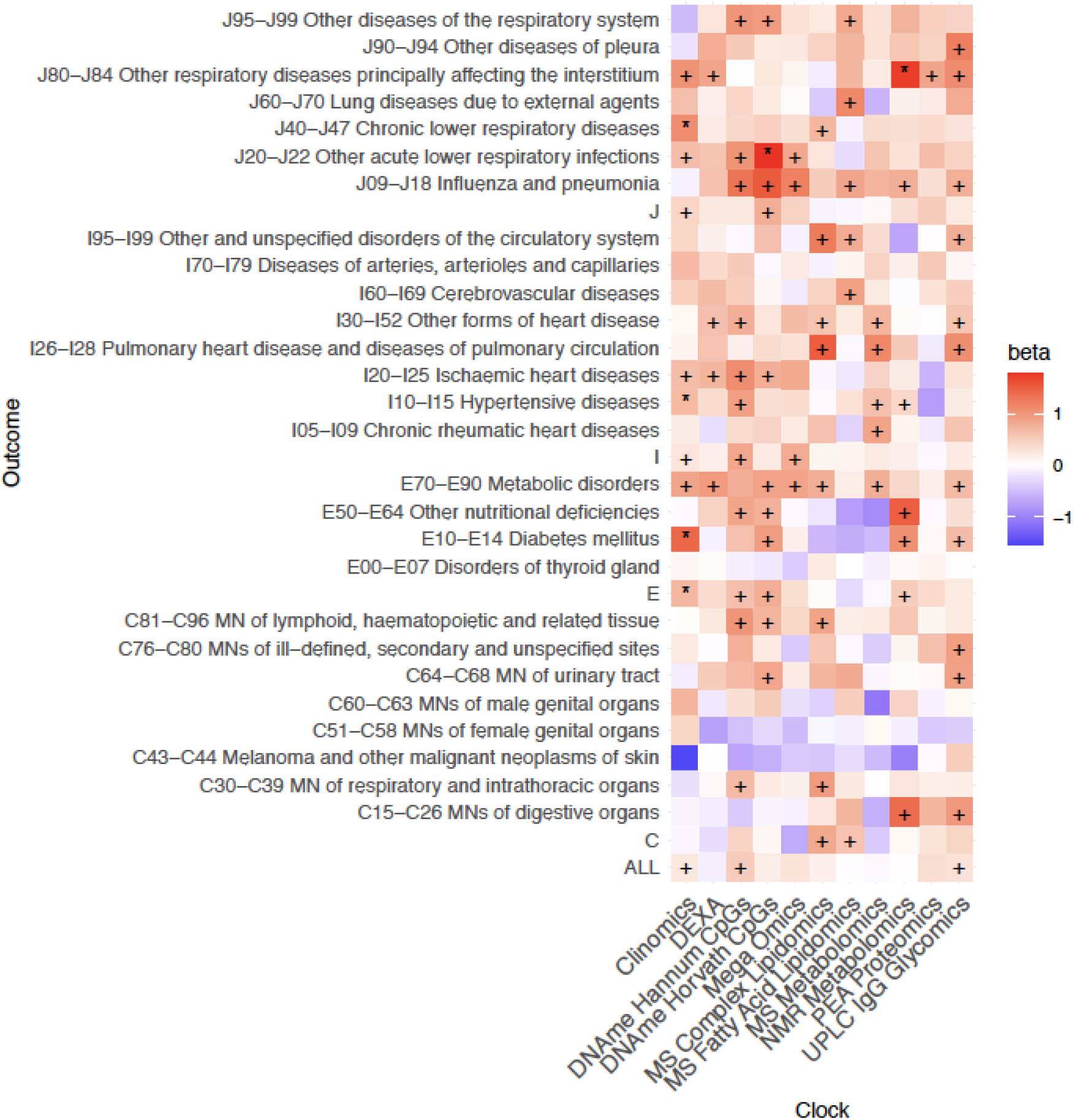
Positive age acceleration associations observed with increased disease risk. Associations with rates of hospitalisation. +/* Association nominally/FDR<10% significant in the frequentist test that OCAA has a positive effect on outcomes. Beta: the relative effect of a year of OCAA to a year of chronAge on disease (initially measured in log_e_ hazard ratios, effect sizes are unitless after division). A value of one indicates that a year of OCAA is equally as deleterious as a year of chronAge and is indicated in salmon colour. To facilitate reading, note the DNAme Horvath CpGs-BMI beta is 1.02 and the DNAme Hannum CpGs-C81-C96I beta is 1.00. Clock: the omics clock on which OCAA was measured. Disease group: the set of diseases (defined by ICD10 codes) which were tested for first incidence after assessment against the clock, already prevalent cases were excluded (Case numbers for each disease block in Supplementary Table 5).

The greater statistical power for risk factors results in considerably more significant associations at FDR<10% (Figure 6). Once more, Clinomics, as might be expected, has the greatest number of significant associations, however NMR metabolomics and UPLC IgG Glycomics OCAAs are nearly as broadly predictive. Mega-omics, MS and NMR Metabolomics OCAAs show positive associations with all risk factors. It should be noted that while the Clinomics OCAA showed most significant FDR<10% associations with diseases and risk factors, its predictors (e.g. cholesterol, FEV1 and SBP) are often close to and designed to predict clinical endpoints and overlap with the risk factors considered here. Similarly, metabolite and lipid-based clocks contain cholesterol subfractions. All OCAAs were associated positively with BMI and total cholesterol. We found strong associations between OCAAs and the marker of inflammation CRP (often with effect sizes >1), meaning OCAA had a larger effect than chronAge. Overall, the averaged effect of OCAA on risk factors as a proportion of the effect on diseases was large for MS Fatty Acid Lipidomics/Clinomics/PEA proteomics (69%/230%/291%) suggesting they are directly tracking the risk factors we considered. Conversely, this proportion was small for Hannum CpGs/Horvath CpGs/UPLC IgG Glycomics (−3%/-1%/29%), suggesting they are prognostic of incident disease and therefore track more generalised ageing (Supplementary Figure 7a).

**Figure 6.**
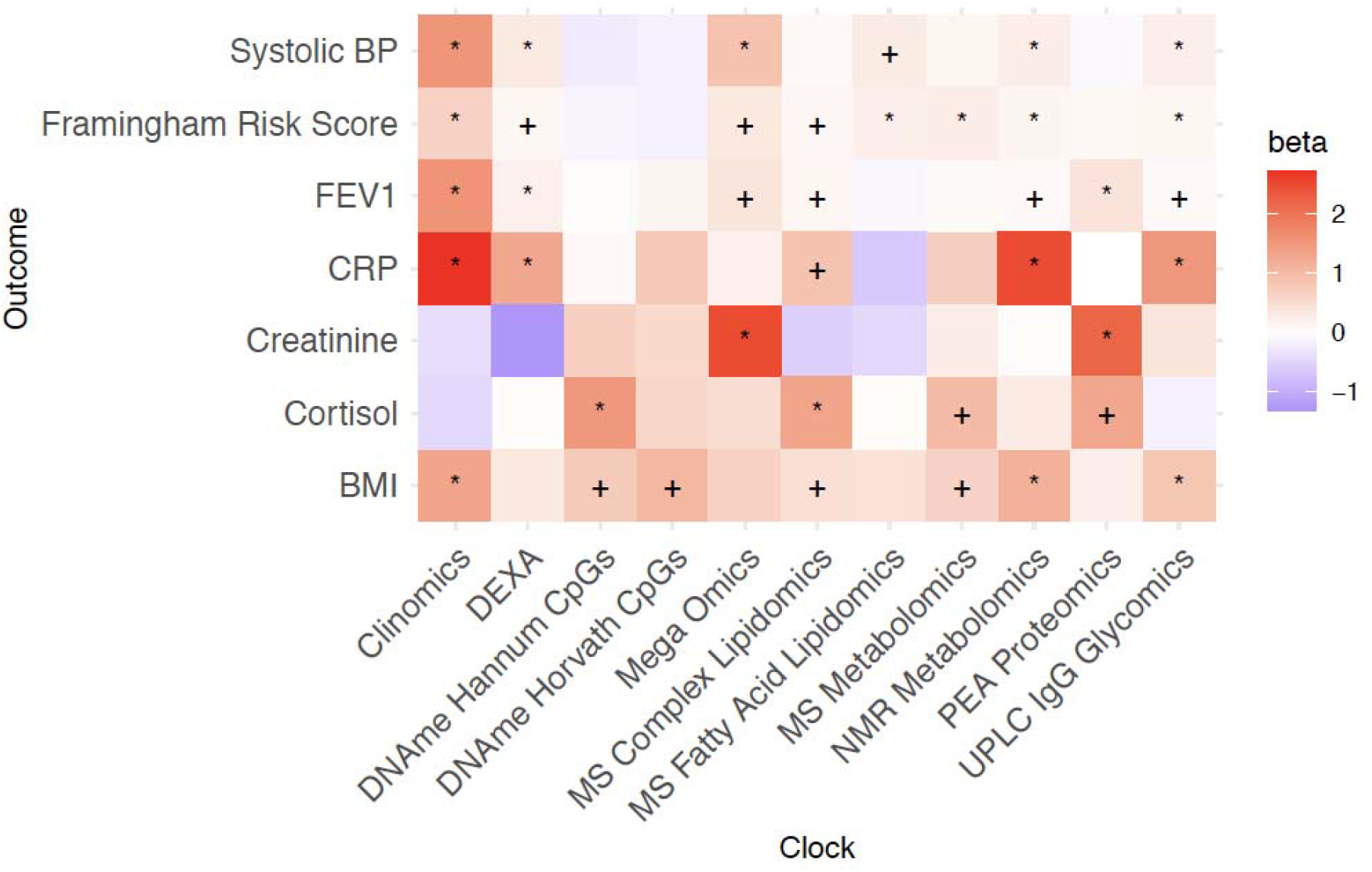
Positive age acceleration associations observed with increased disease risk. Associations with disease risk factors. +/* Association nominally/FDR<10% significant in the frequentist test that OCAA has a positive effect on Risk factors. Beta: the relative effect of a year of OCAA to a year of chronAge on risk factor (effect sizes are unitless after division). A value of one indicates that a year of OCAA is equally as deleterious as a year of chronAge and is indicated in salmon colour. Total cholesterol, which showed a particularly large effect from MS lipidomics OCAA, is excluded here to aid visualisation (the effects on cholesterol can be seen in Supplementary Figure 9b).

We wanted to check if observed OCAA-health associations were driven by the associations of health with smoking and of OCAA with smoking. Our analysis fitting smoking status as a confounder suggests they were not (Supplementary Figures 8a & 8b).

### Comparison of predictive abilities of different OCAAs for risk factors and disease

In principle, two OCAAs could have the same association effect size on disease, but one might be much more prognostic for the population as a whole than the other if it had much larger variation in its range. In order to determine which OCAAs could draw more meaningful distinctions between subjects in terms of health outcomes, we repeated the previous analysis using standardised OCAAs. We found that the standardised Clinomics OCAA showed the greatest predictive power, with an IVW-average effect across all risk factors of 0.39 compared to the range of 0.05-0.12 for the other clocks, with Hannum and Horvath CpGs OCAA smaller still, at −0.014 and 0.018, respectively (SEs ~0.01, in all cases). Conversely, FEV1 was most predictable by standardised OCAA (0.20, SEs ~0.01, IVW-averaged across clocks), whilst creatinine/reversed cortisol were least predictable (0.02/0.04, SEs ~0.01).

Standardised OCAA effects on disease showed an even more uniform pattern (Supplementary Figure 7a): the IVW-average effect across diseases was between 0.11 (MS Metabolomics) and 0.24 (Clinomics), except for the 0.016 of PEA Proteomics (SEs ~0.04). Despite limited power, the disease group showing the most sensitivity to standardised OCAA across clocks was J80-J84 (Other respiratory diseases principally affecting the interstitium; 0.76, SE~0.16), perhaps consistent with the FEV1 finding. Although predictive of risk factors, Clinomics OCAA does not appear unusually predictive of disease. Lung function appears particularly sensitive to both ageing (Supplementary Figure 6b) and OCAA.

### Clocks built from few omics principal components are effective predictors of health outcomes

Finally, we reduced dimensionality and assessed the underlying information about ageing being captured by different omics at the assay level, rather than simply the predictors selected for model inclusion. We constructed clocks using a few principal components (PCs) of omics measures as predictors and repeated the previous analyses with their (standardised) OCAAs, estimating chronAge (Supplementary Figure 10) and predicting health outcomes (Supplementary Figure 11a & 11b). The pattern was striking, the IVW-mean effect sizes across all risk factors of 3 PC OCAAs were more than double our standard OCAAs (Supplementary Figure 11a). For all OCAAs, bar DNAme-based, including more omics PCs in the clocks reduced their ability to estimate distinctions in risk factors. IVW-mean effects on diseases were generally similar for the 3 PC and standard OCAAs, except for the PEA Proteomics OCAA, where 3 PCs-based clock outperformed the standard clock by a factor of 10. Overall, OCAAs derived from a few omic PCs appeared equally predictive as our standard OCAAs for diseases and more predictive for health risk factors.

## Discussion

We have performed the most exhaustive comparison of different omics assays as potential biomarkers of age to date. We have shown firstly, it is possible to construct ageing clocks that produce highly accurate estimations of chronAge with a wide variety of omics biomarkers. Secondly, ageing clocks built using PEA proteomics, DNAme, UPLC IgG glycomics and clinical risk factors in ORCADES were able to estimate chronAge in independent populations. Thirdly, it is possible to achieve the same highly accurate estimation of chronAge using a substantial subset of core biomarkers from each assay. Despite finding only modest positive correlations between our OCAAs, we showed that different clocks overlap in the variation they explain in chronAge more than would be expected by chance if they were independently sampling from a latent set of complete predictors. We found associations of OCAAs with total cholesterol, Framingham Risk Score, C-reactive protein and systolic blood pressure. We found 6 statistically significant (FDR<10%) individual associations and strong evidence of enrichment of association of OCAA with incident disease collectively across our tests (20% were nominally significant p<0.05). We found more variation in OCAA predictiveness across risk factors, than across diseases. Overall, we estimated that one year of OCAA has an effect of 0.46/0.45 years of chronAge on risk factors/disease incidence and showed that OCAA based on clocks built using a few principal components of omics were as prognostic as those presented with all available features.

The correlation of our PEA proteomics, DNAme, UPLC IgG glycomics OCAs and chronAge were similar to published models^5,6,19,20^. Unsurprisingly, DNAme-based clocks built in ORCADES were able to estimate age in both Scottish (Generation Scotland) and Estonian Biobanks (EBB), as the Hannum and Horvath epigenetic clocks have been used successfully in numerous populations. We showed for the first time that clocks built from Olink PEA-based proteomics replicate (in EBB and Croatia-Vis), while clocks built using the SOMAlogic^22^ proteomics platform have been shown to replicate across populations previously. Our UPLC IgG glycomics clock also replicated in an independent population, mirroring the applicability of published GlycanAge measures^19^. Conversely, our NMR metabolomics and DEXA clocks had much lower correlation with chronAge in EBB and UKB. The success of these clocks appears to be study-specific: differences in lifestyle and environmental factors that change with age between the populations of the Orkney Islands and general populations in the UK and Estonia are a plausible cause. This finding serves as a warning as to the generalisability of ageing clocks to new populations.

For a measure of BA to be clinically useful and efficient, effective age estimation based on as few predictors as possible is ideal. We substantially reduced the numbers of biomarkers from each assay that were included in our clocks and showed no dilution of performance across all of our clocks. Enroth *et al*.^20^ showed that this was possible with a protein-based clock, however, we reduced the number of proteins by a larger factor and achieved the same accuracy estimating chronAge. This high performance with a substantial subset of predictors has not previously been shown systematically across nine different types of biomarkers.

The extremely high correlations with chronAge reported, such as the r = 0.97 of the Mega-omics OCA, highlight an issue that has been discussed in prior work: that if enough biomarkers were included in the model it would be possible to perfectly estimate chronAge and, by definition, fail to detect (distinct) BA. Lehallier *et al*.^22^ showed that correlation between OCA and chronAge increases with the number of proteins included in the model. Further, it is possible to explain 100% of the variance in chronAge using DNAme data in large samples^30^. A perfect age predictor would give no information about variation between individuals of the same age and even those which are near perfect will have too little variation in the OCAA to be indicative of health status or outcomes beyond chronAge^31^. We found this trend in our results, that the most accurate estimators of chronAge: Mega-omics, PEA proteomics and MS metabolomics OCAAs were not strongly associated with subsequent hospital admissions, nor DNAme-based OCAAs with risk factors. Of course extremely accurate estimators of chronAge do have their uses, for example in a forensic context^32^, but are not useful in terms of BA. This does not mean the assays themselves cannot be used to estimate BA but highlights a limitation of training ageing clocks on chronAge.

A useful BA must be an indicator of health status or outcomes beyond chronAge. We found DNAme-based OCAAs were better estimators of incident disease than risk factors, consistent with the known performance of Horvath’s epigenetic clock. Several groups have shown Horvath’s DNAme OCAA to be associated with subsequent all-cause mortality^7,33–36^. Differences in Horvath’s OCAA between cases and controls have been found for numerous disease phenotypes^33,37–46^. In contrast, Horvath’s OCAA has been found not to be associated with common risk factors including: LDL cholesterol and CRP^27^, a finding we confirmed. We found that Clinomics and lipid based OCAAs were better at predicting risk factors than disease, whereas the opposite was true for DNAme and UPLC IgG Glycomics OCAAs. The similarity between the predictors in the Clinomics and lipid-based clocks and some of the risk factors could be driving these associations. In contrast, DNAme and UPLC IgG Glycomics being prognostic of incident disease beyond chronAge suggests they are more likely to be capturing underlying BA.

It is perhaps not surprising that the Clinomics OCAA showed the strongest evidence of association with disease - it used common clinical measures thought to be prognostic of health. Nonetheless, the pattern is a reassuring proof of concept. The overall enrichment of OCAA-disease and -risk factor associations, strengthens the case for the notion of BA, trackable through omics markers. Jansen *et al*.^9^ showed that their NMR Metabolomics OCAA was significantly higher in cases of metabolic syndrome and cardiometabolic disease than controls, however was not prognostic of incident disease. Our NMR Metabolomics OCAA was nominally associated (p<0.05) of several metabolic disease blocks, suggesting that in a more powered sample this relationship would be significant. Previously, it has been shown that GlycanAge is associated with risk factors^19^ and that IgG glycans (i.e not an OCAA, rather the glycan levels themselves) are effective predictors of incident type 2 diabetes and cardiovascular events^47–49^. However, we are the first to show UPLC IgG glycomics OCAA to be prognostic of incident disease and highlight this is not simply due to tracking the risk factors we considered.

As by definition, having a BA of +1 indicates that the individual has the same functional capacity and risk of age-related disease as the average individual that is one calendar year older than them, indicating the effect of true BA is the same as 1 year of chronAge. Our estimate that the mean effect of 1 year of OCAA on disease incidence is the same as 0.45 years of chronAge is important. BA thus appears to be real and measurable and have effects of similar magnitude to chronAge, albeit our estimates are significantly diluted compared to chronAge, possibly due to OCAA capturing only some aspects of BA, reflecting the types of assay and tissue, rather than BA itself. Better measures of BA seem worthy of pursuit, as do interventions that can reverse well-measured BA. The negative association between Melanoma and other malignant neoplasms of skin (C43-C44) and OCAAs for many clocks leads us to suggest a less sedentary lifestyle is leading to lower OCAA, but also increased exposure to the sun. If replicated, this will highlight that skin BA and other BAs need not closely align, and we speculate this finding might also generalise across other organs.

A strength of our work was the sheer number and range of assays and therefore omics ageing clocks whose performance we compared in the same individuals, whereas previous comparisons have been limited to DNAme-based clocks^25,50,51^ or DNAme, clinical risk factors and frailty measures^52^. We have tried to validate our omics ageing clocks trained in ORCADES in independent populations where available, to illustrate their wider applicability. A limitation faced by previous studies was the narrow age range of individuals in the training sample, for example Lee *et al*.’s epigenetic clock trained in a pregnancy cohort produced extremely accurate estimations of chronAge for individuals under 45 but underestimated age in older individuals^53^. Our clocks avoid this limitation due to the wide age range (16-100) of individuals in the ORCADES cohort.

The novel assessment of excess overlap between clocks is a strength of this work, as it has not previously been shown that, across multiple different omics assays, OCAs overlap more than would be expected by chance if they were ISLSP, indicating these clocks are tracking more common rather complementary aspects of ageing. A further strength is the regularisation of effect sizes - we have measured the effect of OCAA per effect of year of chronAge - giving a natural and understandable scale. Another strength is its scope, with many clocks tested against many age-related diseases. Of course, this is also a weakness, as it reduces power after compensation for multiple testing. Nonetheless, the essentially agnostic view taken of individual disease groupings and clocks does mitigate the risk of publication bias.

A limitation of this work is the relatively small sample size, both in terms of the number of individuals with multiple omics assays and within that, the number of incident hospital admissions over the follow-up period. Due to the low number of deaths in our sample we are as yet unable to test for the association of OCAA on mortality, as in previous studies. As the omics data available for ORCADES is cross-sectional, we were unable to comment on the variation of OCAA within individuals over time. However, we were able to investigate the prognostic ability of single time point OCAAs on hospital admissions over a 10-year follow up. The nature of our sample, a population isolate, means there is potential for local factors to influence our results. We have shown this is not the case for several of our omics clocks’ accuracies (Supplementary Figure 2), as they were successfully replicated in additional populations, however, it could contribute to the poor replication seen for the DEXA and NMR metabolomics clocks. The use of hospitalisation as a measure of incidence is a limitation, particularly acute for diseases normally treated in the community such as type 2 diabetes and influenza. Nonetheless, we are likely to have captured the most severe cases and have tested whether this severity associates with OCAA and presumed frailty, giving rise to more severe experience of the disease. Secondly, the correlated nature of the assays and of the disease outcomes mean our tests have not been independent, although this means the FDR corrections have been conservative. A more powered study might also try to disentangle individual markers especially those retained in our core omics clocks and consider their biological plausibility as sitting on the causal pathway.

Of course, association does not imply causation. Although the use of a prospective cohort has reduced the risk of reverse causation, undiagnosed cases (at baseline) might still have contributed to the effects we observe, although confounding where a latent set of underlying traits is influencing disease susceptibility and the biomarkers is perhaps more likely. Nonetheless, even in the absence of causation, OCAA does appear to often be a biomarker of disease and underlying BA.

In conclusion, our work has strongly further evidenced the existence of BA as distinct from chronAge, whilst highlighting a substantial part of the OCAA is noise. The data also suggested there may be more than one type of BA, as measured by different clocks and giving rise to differing amounts of disease susceptibility, most strongly implied by our evidence that skin age and heart age may move in opposite directions. We also highlight that some OCAAs (e.g. PEA proteomics) may capture specific risks and consequently associate with health, whilst others (e.g. DNAme and UPLC IgG glycomics) may capture more generalised ageing. Our observation that clocks derived from few PCs of omics are less accurate in estimating chronAge but better able to predict risk factors, suggests that the search for BA should be pursued through salient features of biology. This supports the recent success of ageing clocks trained on all-cause mortality based measures^2,54^, DNAmePhenoAge^2^ and GrimAge^54^, which have been shown to be more prognostic of health and mortality outcomes than DNAme clocks trained on chronAge directly^24,25,52,55^. Similarly, the mortality trained NMR Metabolomics measure from Deelen *et al*. is more prognostic of both 5- and 10-year all-cause mortality than a model of conventional mortality risk factors^18^. We therefore suggest that the focus of future research should continue to shift to clocks trained on mortality, or more ideally all-cause morbidity, that are prognostic of subsequent health outcomes rather than accurate chronAge estimators.

## Acknowledgements

We thank Archie Campbell for assistance with the electronic health records from ORCADES. The Orkney Complex Disease Study (ORCADES) was supported by the Chief Scientist Office of the Scottish Government (CZB/4/276, CZB/4/710), a Royal Society URF to J.F.W., the MRC Human Genetics Unit quinquennial programme “QTL in Health and Disease”, Arthritis Research UK and the European Union framework program 6 EUROSPAN project (contract no. LSHG-CT-2006-018947). DNA extractions were performed at the Edinburgh Clinical Research Facility, University of Edinburgh. We would like to acknowledge the invaluable contributions of the research nurses in Orkney, the administrative team in Edinburgh and the people of Orkney. We acknowledge support from the MRC Human Genetics Unit programme grant, “Quantitative traits in health and disease” (U. MC_UU_00007/10).

The CROATIA-Vis and, CROATIA-Korcula studies were funded by grants from the Medical Research Council (UK), from the Republic of Croatia Ministry of Science, Education and Sports (108-1080315-0302; 216-1080315-0302) and the Croatian Science Foundation (8875); and the CROATIA-Korcula genotyping was funded by the European Union framework program 6 project EUROSPAN (LSHGCT2006018947). We thank the staff of several institutions in Croatia that supported the field work, including Zagreb Medical Schools, the Institute for Anthropological Research in Zagreb, the recruitment team from the Croatian Centre for Global Health, University of Split and all the study participants. We are grateful to the Helmholtz Zentrum Munchen (Munich, Germany), AROS Applied Biotechnology, (Aarhus, Denmark) and the Edinburgh Clinical Research facility, University of Edinburgh (Edinburgh, United Kingdom) for SNP array genotyping.

Generation Scotland received core support from the Chief Scientist Office of the Scottish Government Health Directorates [CZD/16/6] and the Scottish Funding Council [HR03006] and is currently supported by the Wellcome Trust [216767/Z/19/Z]. Genotyping of the GS:SFHS samples was carried out by the Genetics Core Laboratory at the Edinburgh Clinical Research Facility, University of Edinburgh, Scotland and was funded by the Medical Research Council UK and the Wellcome Trust (Wellcome Trust Strategic Award “STratifying Resilience and Depression Longitudinally” (STRADL) Reference 104036/Z/14/Z). We are grateful to all the families who took part, the general practitioners and the Scottish School of Primary Care for their help in recruiting them, and the whole Generation Scotland team, which includes interviewers, computer and laboratory technicians, clerical workers, research scientists, volunteers, managers, receptionists, healthcare assistants and nurses. Ethical approval for the GS:SFHS study was obtained from the Tayside Committee on Medical Research Ethics (on behalf of the National Health Service).

This research was funded by the European Union through the European Regional Development Fund (N.T.) and SP1GI18045T (T.E., N.T.), grants PUT1660 (T.E., N.T.) and PUT1665 (K.F., N.T.) of the Estonian Research Council. The Estonian Biobank study was approved by the Ethics Review Committee on Human Research of the University of Tartu and the Estonian Genome Center, University of Tartu scientific committee. Written informed consent was obtained from participants in accordance with the Declaration of Helsinki. The work of L.K. was supported by an RCUK Innovation Fellowship from the National Productivity Investment Fund (MR/R026408/1).

We thank the UK Biobank Resource, approved under application 19655. We acknowledge funding from the UK Medical Research Council Doctoral Training Programme in Precision Medicine (MR/N013166/1).

## Author Contributions

E.M.D: Project conception, methodology, manuscript drafting and editing, formal analysis, visualisation. N.T: Performed validation analysis in Estonian Biobank. L.K: Processed and performed QC on the UPLC IgG Glycomcis data in ORCADES, Croatia-Vis & Croatia-Korcula and supervised A.F. A.F: Processed and performed QC on the DNA methylation data in ORCADES. R.W: Preparation of the DNA methylation data in GS:SFHS. P.K.J: Project conception, methodology, formal analysis, manuscript drafting and editing, project supervision. J.F.W: Project conception, manuscript drafting and editing, project supervision. All other authors commented and approved the manuscript prior to submission.

## Competing Interests

P.K.J is a paid consultant for Humanity Inc. and Global Gene Corporation.

## Data Availability

There is neither research ethics committee approval, nor consent from individual participants, to permit open release of the individual level research data underlying this study. Please contact the QTL Data Access Committee for further information if required. Access to data from GS:SFHS is by application and a managed process. Access to data from the Estonian Biobank is by request and managed by the Estonian Committee on Bioethics and Human Research.

## Code Availability

## Methods

### Cohort Data

Analyses were predominantly carried out using the Orkney Complex Disease Study (ORCADES)^28^, a population-based isolate cohort that is extensively characterised in terms of both traditional phenotypes, omics assays and mean 12 years of follow up via linked electronic health records (EHR). The additional cohorts, Croatia-Vis and Croatia-Korčula^56,57^, were used to validate omics ageing clocks trained in ORCADES. Croatia-Vis was used to validate a clock trained in ORCADES using a subset of proteins (those measured on the Olink CVDII, CVDIII and INFI panels) referred to as protein subset 1 and the UPLC IgG glycomics clock. Replication of the NMR metabolomics and UPLC IgG glycomics clocks trained in ORCADES was carried out in Croatia-Korčula. The Estonian Biobank^58^ (EBB) cohort was used to validate a clock trained using a subset of proteins (those measured on the Olink CVII, CVDIII, INF1 and ONCII panels) referred to as protein subset 2 as well as the NMR Metabolomics clock. Both EBB and the Generation Scotland: Scottish Family Health Study (GS:SFHS)^59^, a family-based cohort comprising volunteers across Scotland, were used to assess two DNAme-based ageing clocks. Finally, the UK Biobank^60^ (UKB) was used to test the Clinomics and DEXA clocks trained in ORCADES.

### Omics Assays

#### Dual X-ray absorptiometry (DEXA)

Whole body imaging was performed on the Hologic fan beam DEXA scanner (GE Healthcare). Measures of body composition were derived from the DEXA scans using APEX2 software for bone, lean and fat tissue and APEX4 software for android, gynoid, visceral and lean fat mass content. 28 measures in the following broad categories: bone mineral density, bone mineral content, fat or lean mass percentages for head, trunk and limbs were selected for analyses. These were measures that did not use chronAge in their calculation and were also available in the UK Biobank. Measures were removed as outliers based on a z-score cut-off of 6 then pre-corrected for sex. Residuals were additionally subject to a threshold by removing outliers with a z-score cut-off of 3.

#### DNA Methylation

The Illumina EPIC 850K array was used to measure DNA methylation levels in ORCADES. Quality control was carried out using the meffilQC pipeline^61^ and minfi package^62^. Samples were excluded as outliers if >1% of probes had a detection p-value > 0.01, due to failure of sex concordance, if samples showed evidence of dye bias or failed median methylation signal z-score cut-off of 3. Probes were removed as outliers if the detection p-value was >0.01 in >1% of samples or had a bead count of <3 in at least 5% of samples. The *preprocessNoob* function in the “minfi” package was used for array normalisation to remove unwanted technical variation. M values were corrected for the technical covariates: plate number (as a random effect), season of venepuncture, year of venepuncture, plate position and 10 principal components of the control probes (as fixed effects) using GCTA-REML^63^.

Instead of creating novel DNA methylation clocks when there are landmark clocks available in the literature, we constructed clocks based on Hannum and Horvath’s original epigenetic clocks, to compare with our other omics. As ORCADES used the Illumina EPIC 850k chip rather than the earlier 450k/27k chips used by Hannum and Horvath, our methylation clocks are subsets of Hannum and Horvath’s clocks. It has been shown that imputing probes that are absent from the 850k chip but present in the 450k/27k set leads to underestimation of both published ageing measures^64^. Thus, for our clocks named Hannum CpGs and Horvath CpGs we presented 62/71 and 333/353 of sites, respectively, that were present on the 850k chip to the penalised regression algorithm for model selection. Residuals from REML within a z-score threshold of 6 were then corrected for sex.

#### NMR Metabolomics

The high throughput NMR metabolomics assay of EDTA plasma (Nightingale Health Ltd., Helsinki, Finland) quantified 225 metabolomics measures in molar concentration units. The measures include amino acids, ketone bodies, low molecular weight metabolites and numerous lipid and lipoproteins subclasses. In both ORCADES and Croatia-Korčula, metabolite measures were removed as outliers based on a z-score cut-off of 6, pre-corrected for sex and the use of statins as a binary variable. Residuals were additionally removed as outliers with a z-score cut-off of 3.

#### MS Fatty Acids Lipidomics

Shotgun lipidomics and liquid chromatography tandem mass spectrometry (LC-MS/MS) was used to quantify the molar concentrations of 44 fatty acids as described previously^65^. Fatty acid measures were removed as outliers based on a z-score cut-off of 6, pre-corrected for sex, box number, plate position and use of statins.

#### UPLC IgG Glycomics

The glycan data have previously been described in detail by Kristic *et al*., for the ORCADES^19^, Croatia-Vis and Croatia-Korcula^56,57^ studies. Raw glycan measures were total area normalised and batch corrected using the “ComBat” function of the sva package^66^ in R. The normalised glycan measures were excluded as outliers based on a z-score threshold of 6 and pre-corrected for sex.

#### PEA Proteomics

1,102 proteins were measured using a proximity extension assay method (Olink Bioscience, Uppsala, Sweden)^67^ from EDTA plasma in 12 x 92-protein panels designated by the manufacturer: cardiovascular 2, cardiovascular 3, inflammation 1, metabolism, cardiometabolic, cell regulation, development, immune response, organ damage, oncology 2, neurology and neuro-exploratory. Measures for all twelve panels are available for 1,057 individuals in ORCADES, with subsets available in Croatia-Vis (inflammation 1, cardiovascular 2 and cardiovascular 3) and EBB (inflammation 1, cardiovascular 2, cardiovascular 3 and oncology 2). PEA proteomics-based OCAs were rederived using these subsets to allow comparison across populations. NPX values of proteins (on the log2 scale) including those non-missing below the lower limit of detection (LOD), were removed as outliers with a z-score cut-off of 6. These measures were then precorrected for the following covariates via fixed effects linear regression: sex, season of venepuncture, time the plasma sample was in storage between collection and assay (days), plate number, plate row and plate column.

#### Clinomics

This dataset consisted of 13 selected clinical measures that are routinely measured during visits with general practitioners and clinicians: albumin, fasting plasma glucose, calcium, uric acid, high density lipoprotein cholesterol, total cholesterol, triglycerides, height, weight, forced expiratory volume in 1 second (FEV1), and diastolic and systolic blood pressure.

#### MS Metabolomics & MS Complex Lipidomics

Non-targeted metabolomic and lipidomic features were detected and quantified using Metabolon as described previously^68^. The HD4 dataset comprised measures of 1143 biochemicals while the complex lipids dataset measured 1052 biochemicals, these were treated as two separate omics assays referred to as MS Metabolomics and MS Complex Lipidomics respectively. Measures were removed as outliers with a z-score cut-off of 6. These measures were then pre-corrected for the following covariates via fixed effects linear regression: sex, statin use, assay run day, plate number and plate row and plate column.

#### EHR

The ORCADES cohort has record linkage to hospital admission records (Scottish Morbidity Records: SMR01). The first occurrence of any hospital admission with ICD10 diagnosis, was taken as incidence. NHS Scotland records moved from ICD9 to ICD10 in April 1996, so diagnoses since ~12 years prior to assessment were captured. The disease groupings analysed included each ICD10 block within 5 Chapters thought a priori to associate with age II (Neoplasms - codes C), IV (Endocrine, nutritional and metabolic diseases - codes E), IX (Diseases of the circulatory system - codes I), and X (Diseases of the respiratory system - codes J). For Chapter II only C codes (malignant) were included in our analyses. Chapters as a whole were also analysed, as were all the diseases from these chapters simultaneously. Incident disease was defined as the time of first hospital admission with a diagnostic code recorded (in any position in the admission record) for any disease within the grouping being analysed. For each disease grouping, subjects with recorded admission prior to the date of venepuncture were then excluded entirely in the subsequent analysis, as already prevalent.

### Quality Control of Omics Measures

Outliers were defined based on z-score thresholds that varied between omics datasets depending on the distributions of the raw measures. Omics measures were pre-corrected for known batch effects and covariates (specified above) using fixed effects linear regression or other specified methods. A second pass z-score threshold on the residuals was used to detect further outliers for a subset of assays and all missing values were removed. The residuals produced from covariate correction were then scaled and centred to have a mean of zero and a standard deviation of one to ensure that effect sizes of any variables included in the models were comparable.

### Clock Construction

#### Per Omics Assay

The individuals in the ORCADES cohort were split into 75% training, 25% testing. For the analysis comparing clock performance across omics platforms the testing 25% of samples were taken preferentially from the pool of individuals that possess measures for all of the omics platforms. Tenfold cross validation in the training sample was used to select the shrinkage parameter, λ, for the penalised regression that was estimated to produce the model with the minimum mean squared error. Models were constructed using three different procedures implemented using the glmnet^69^ and caret packages in R with chronAge at venepuncture as the dependent variable: i) least absolute shrinkage and selection operator (LASSO) regression ii) elastic net regression with an alpha of 0.5 iii) elastic net regression with alpha select using 10-fold cross validation in the training sample. We found no difference in performance between the three methods so construction using elastic net regression with an alpha of 0.5 was used throughout the analyses presented. This model was then used to estimate chronAge in the testing sample and an independent out of cohort sample if available.

As stochasticity is present in the procedure, the variables selected for model inclusion will vary depending on the individuals selected to be in the training sample, clock construction was repeated 500 times and the features selected for inclusion and the correlation between chronAge and age estimated by the model were recorded to ensure that the model performance results presented here are representative and not an outlier due to individuals at extreme ends of distributions contributing to the training sample and rare model being used to draw conclusions (data not shown).

#### Mega-Omics

model that was presented with all of the features from all of the omics platforms. The dataset itself was created by merging all of the corrected omics measures (residuals) after platform level quality control, again standardising all features to have a mean of zero and standard deviation of one. The clock was created using the same construction procedure outlined above.

#### Core Models

models were constructed per omics assay. The elastic net regression algorithm was presented with only those predictors that were selected for model inclusion in >95% of the 500 iterations of clock construction for the relevant omics platform. This reduced set of predictors then underwent clock construction as described above.

#### Principal Component Clocks

To ensure that the differences in variance explained in chronAge by different omics clocks is not due to the discrepancy between the number of features available and hence the number of features selected for model inclusion across omics types but rather a genuine difference in the information about ageing captured by different omics; clocks were built using principal components (PCs) of the relevant omics platform as features. The first 3, 5, 10 and 20 PCs were extracted from the covariate corrected scaled and centred omics data at the platform level using the *prcomp* function in R. These PCs were then presented to the elastic net algorithm and clocks built.

### Correlation of OCAAs

Pairwise Pearson correlations between 10 of our OCAAs were calculated, Mega-omics OCAA was excluded from this and all between clock comparisons as it contains predictors spanning multiple assays.

### Partitioning Variance Explained in ChronAge

The unique variance in chronAge explained by each clock, 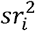, was calculated as the squared part correlation of chronAge (*Y*) and age estimated by clock *i* while controlling for all of the other *k* clocks. Part correlations were calculated using the *spcor.test* function in the “ppcor” package in R^70^. The portion of variance in chronAge explained by all of the *k* clocks together, the *R*^2^ from the following model:

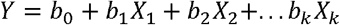

Where *Y* is chronAge and *X*_1…k_ are age estimated by clocks 1 to *k*, was used to partition the total variance of chronAge further into that which remains unexplained by the 10 clocks (1 - *R*^2^) and that which is explained by overlapping clocks:

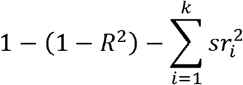

To gain a more detailed insight into the relationship between clocks we carried out pairwise comparisons. Following the same procedure as outlined above, the unique variance in chronAge explained by each clock in the pair is the squared part correlation of chronAge and age estimated by one clock while controlling for age estimated by the other clock in the pair. The variance remaining unexplained by either of the clocks was 1 - *R*^2^ of a bivariate model. The overlap, calculated by subtraction, is specifically the variance in chronAge explained by both of the clocks in the pair. This is unlike overlap calculated in the previous step, where we were only able to state that this variance was not unique to a particular clock but unable to deconstruct further.

### Assessing the Overlap between Clocks

We assessed whether the combined variance in chronAge explained by pairs of clocks deviated from what would be expected by chance if both clocks were independently sampling from a latent set predictors (ISLSP) of chronAge. The combined variance in chronAge explained by both clocks together was calculated as the multiple *R*^2^ from a bivariate model, with chronAge being the dependent variable and the estimated ages from the two clocks in the pair the independent variables. The variance explained in chronAge (*v*_i_) by each clock (*i*) individually was the univariate *R*^2^ from the regression of estimated age on chronAge. The expected variance in chronAge explained by two clocks by chance (*E*) was calculated as follows:

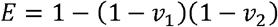

The idea being that the variance in chronAge not already explained by the first clock is 1 - *V*_i_. With the null hypothesis that the two clocks are independent samples from the latent set of complete predictors and thus explain partly overlapping information about age. The expected left unexplained after the addition of the second clock is thus (1 - *v*_1_)(1 - *v*_2_).

To allow for the comparison of the deviation of observed variance explained in chronAge (*O*) from expected (*E*) across pairs of clocks, this deviation was re-scaled. As the magnitude of *v*_i_ effects the possible range of values *O* could take. The theoretical minimum variance explained (*E*_min_) by two clocks is the variance explained by the larger of the two clocks alone (the second clock only providing information already captured by the first). The theoretical maximum (*E*_max_) is *v*_1_ + *v*_2_ or 1 if *v*_1_ + *v*_2_ > 1 (the clocks are explaining entirely nonoverlapping variance). Comparisons containing clocks with high *v*_i_ will have a much smaller range of possible *0* than those with low *v*_i_ so directly comparing the magnitude of the deviation of observed from expected is not ideal. The results presented are on a scale of excess overlap calculated as follows:

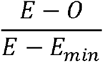

With a value of 0 meaning that the observed variance explained equals that expected by chance if the clocks were independent. A value of 1 denoting that no additional variance was explained with the addition of the second clock. Negative values are possible and mean that the two clocks overlap less than expected and track separate aspects of chronological ageing, but in practice, we see that the clocks always track more common aspects than would be expected under the null hypothesis, albeit to varying degrees.

### Association with health-related phenotypes & Incident Disease

OCAAs were tested for association with health-related risk factors and age-related incident diseases, as measured by hospital admission.

#### Association with chronAge

We first tested whether the risk factors and disease outcomes were associated with chronAge. For incident disease: time from assessment to incidence or to study end (the date when SMR01 records were extracted: December 2017, around ten years after assessment) was modelled using a Cox proportional hazard model^71^ and the Surv function in the “survival” package in R. Subjects with prevalent disease were excluded. The baseline hazard was dependent on time since assessment, and hazards ratios dependent on chronAge and sex. We used time since assessment as the determinant of base hazard rather than chronAge, so that we could determine which groupings had stronger age-related effects and compare the effects of OCAA to those of chronAge. P-values for association with chronAge (and later OCAA) were calculated using a one-sided test, with H1 being that chronAge increased risk.

#### Association with OCAAs

with standardised risk factors (units of phenotypic standard deviation) were carried out using linear regression with chronAge and sex fitted as fixed effects covariates. To restrict the burden of multiple testing we only tested the association of OCAAs on risk factors or disease blocks which showed a statistically significant association (effect size >0) with chronAge at outset (Benjamini-Hochberg FDR<10%) and had >5 incident cases (disease blocks). We tested the effect of OCAAs on each disease grouping using the same model as for chronAge, including chronAge and OCAA as effects. OCAA was not standardised but observed effect sizes were rescaled (divided) by the effect of chronAge, using the same model, enabling a comparison of the effect of one year’s OCAA with one year’s chronAge, with a value of 1 denoting the same effect. False discovery rate was again determined using the Benjamini-Hochberg method (FDR<10%).

Across both risk factors and disease, we found that large estimated effects arose in the context of large SEs. To facilitate visualising the results we had most confidence in we applied a shrinkage method, imposing a prior assumption on the distribution of beta (mean 0, SD 1) to the likelihood of our observed beta, shrinking resultant estimates with larger SEs more towards 0.

Individual tests of association generally had limited power due to multiple testing and the low variance of OCAA (compared with chronAge). We therefore considered the results of each OCAA across multiple outcomes by inverse variance weighting (IVW) observed results for individual outcomes. The covariance amongst outcomes and predictors, mean that the independence assumption for meta-analysis (or sign testing) is violated. Whilst this should not bias estimates, their precision will be overstated. We consider these results to be descriptive, and not conformable to formal testing. We use “~” to denote SEs calculated under the violated independence assumption, but still consider these useful to give a sense of magnitude. Conversely, for the same reason, the formal tests we perform (FDRs) are likely to be conservative.

We repeated these analyses with standardised OCAAs to compare the prognostic ability of different OCAAs at a population level, across risk factors and diseases and with our PC clocks OCAAs.

## References

1. López-Otín, C., Blasco, M. A., Partridge, L., Serrano, M. & Kroemer, G. The Hallmarks of Aging. Cell 153, 1194–1217 (2013).

2. Levine, M. E. et al. An epigenetic biomarker of aging for lifespan and healthspan. Aging 10, 573–591 (2018).

3. Horvath, S. et al. Reversing age: dual species measurement of epigenetic age with a single clock. doi:10.1101/2020.05.07.082917.

4. Baker, G. T. & Sprott, R. L. Biomarkers of aging. Exp. Gerontol. 23, 223–239 (1988).

5. Hannum, G. et al. Genome-wide Methylation Profiles Reveal Quantitative Views of Human Aging Rates. Mol. Cell 49, 359–367 (2013).

6. Horvath, S. DNA methylation age of human tissues and cell types. Genome Biol. 14, R115–R115 (2013).

7. Marioni, R. E. et al. DNA methylation age of blood predicts all-cause mortality in later life. Genome Biol. 16, 25 (2015).

8. Bocklandt, S. et al. Epigenetic Predictor of Age. PLOS ONE 6, e14821 (2011).

9. Jansen, R. et al. An integrative study of five biological clocks in somatic and mental health. eLife 10, e59479 (2021).

10. Zhang, W. G. et al. Select aging biomarkers based on telomere length and chronological age to build a biological age equation. Age 36, 1201–1211 (2014).

11. Chen, W. et al. Three-dimensional human facial morphologies as robust aging markers. Cell Res. 25, 574–587 (2015).

12. Cole, J. H. et al. Predicting brain age with deep learning from raw imaging data results in a reliable and heritable biomarker. NeuroImage 163, 115–124 (2017).

13. Cole, J. H. et al. Brain age predicts mortality. Mol. Psychiatry 23, 1385–1392 (2018).

14. Goyal, M. S. et al. Persistent metabolic youth in the aging female brain. Proc. Natl. Acad. Sci. 116, 3251 (2019).

15. Sokolova, K., Barker, G. J. & Montana, G. Convolutional neural-network-based ordinal regression for brain age prediction from MRI scans. in Medical Imaging 2020: Image Processing vol. 11313 113132B (International Society for Optics and Photonics, 2020).

16. Fischer, K. et al. Biomarker Profiling by Nuclear Magnetic Resonance Spectroscopy for the Prediction of All-Cause Mortality: An Observational Study of 17,345 Persons. PLoS Med. 11, e1001606–e1001606 (2014).

17. Menni, C. et al. Metabolomic markers reveal novel pathways of ageing and early development in human populations. Int. J. Epidemiol. 42, 1111–1119 (2013).

18. Deelen, J. et al. A metabolic profile of all-cause mortality risk identified in an observational study of 44,168 individuals. Nat. Commun. 10, 3346–3346 (2019).

19. Krištić, J. et al. Glycans Are a Novel Biomarker of Chronological and Biological Ages. J. Gerontol. Ser. A 69, 779–789 (2014).

20. Enroth, S., Enroth, S. B., Johansson, Å. & Gyllensten, U. Protein profiling reveals consequences of lifestyle choices on predicted biological aging. Sci. Rep. 5, 1–10 (2015).

21. Lehallier, B. et al. Undulating changes in human plasma proteome across lifespan are linked to disease. bioRxiv 751115–751115 (2019) doi:10.1101/751115.

22. Lehallier, B., Shokhirev, M. N., Wyss-Coray, T. & Johnson, A. A. Data mining of human plasma proteins generates a multitude of highly predictive aging clocks that reflect different aspects of aging. Aging Cell n/a, e13256.

23. Alpert, A. et al. A clinically meaningful metric of immune age derived from highdimensional longitudinal monitoring. Nat. Med. 25, (2019).

24. Hillary, R. F. et al. Epigenetic measures of ageing predict the prevalence and incidence of leading causes of death and disease burden. Clin. Epigenetics 12, 115 (2020).

25. Maddock, J. et al. DNA methylation age and physical and cognitive ageing. J. Gerontol. Ser. A (2019) doi:10.1093/gerona/glz246.

26. Zheng, Y. et al. Blood Epigenetic Age may Predict Cancer Incidence and Mortality. EBioMedicine 5, 68–73 (2016).

27. Horvath, S. et al. An epigenetic clock analysis of race/ethnicity, sex, and coronary heart disease. Genome Biol. 17, 171 (2016).

28. McQuillan, R. et al. Runs of Homozygosity in European Populations. Am. J. Hum. Genet. 83, 359–372 (2008).

29. Franke, K., Ziegler, G., Klöppel, S. & Gaser, C. Estimating the age of healthy subjects from T1-weighted MRI scans using kernel methods: Exploring the influence of various parameters. NeuroImage 50, 883–892 (2010).

30. Zhang, Q. et al. Improved precision of epigenetic clock estimates across tissues and its implication for biological ageing. Genome Med. 11, 54–54 (2019).

31. Pyrkov, T. V. & Fedichev, P. O. Biological age is a universal marker of aging, stress, and frailty. doi:10.1101/578245.

32. Alsaleh, H. & Haddrill, P. R. Identifying blood-specific age-related DNA methylation markers on the Illumina MethylationEPIC^®^ BeadChip. Forensic Sci. Int. 303, 109944 (2019).

33. Horvath, S. et al. Decreased epigenetic age of PBMCs from Italian semi-supercentenarians and their offspring. Aging 7, 1159–1170 (2015).

34. Hofman, A. et al. DNA methylation-based measures of biological age: meta-analysis predicting time to death. Aging 8, 1844–1865 (2016).

35. Christiansen, L. et al. DNA methylation age is associated with mortality in a longitudinal Danish twin study. Aging Cell 15, 149–154 (2016).

36. Perna, L. et al. Epigenetic age acceleration predicts cancer, cardiovascular, and allcause mortality in a German case cohort. Clin. Epigenetics 8, 64 (2016).

37. Levine, M. E., Lu, A. T., Bennett, D. A. & Horvath, S. Epigenetic age of the pre-frontal cortex is associated with neuritic plaques, amyloid load, and Alzheimer’s disease related cognitive functioning. Aging vol. 7 1198–1211 https://www.aging-us.com/article/100864/text (2015).

38. Vidal, L. et al. Specific increase of methylation age in osteoarthritis cartilage. Osteoarthritis Cartilage 24, S63 (2016).

39. Marioni, R. E. et al. The epigenetic clock is correlated with physical and cognitive fitness in the Lothian Birth Cohort 1936. Int. J. Epidemiol. 44, 1388–1396 (2015).

40. Horvath, S. & Ritz, B. R. Increased epigenetic age and granulocyte counts in the blood of Parkinson’s disease patients. Aging vol. 7 1130–1142 https://www.aging-us.com/article/100859/text (2015).

41. Maierhofer, A. et al. Accelerated epigenetic aging in Werner syndrome. Aging vol. 9 1143–1152 https://www.aging-us.com/article/101217/text (2017).

42. Horvath, S. et al. Huntington’s disease accelerates epigenetic aging of human brain and disrupts DNA methylation levels. Aging 8, 1485–1504 (2016).

43. Breitling, L. P. et al. Frailty is associated with the epigenetic clock but not with telomere length in a German cohort. Clin. Epigenetics 8, 21 (2016).

44. Walker, R. F. et al. Epigenetic age analysis of children who seem to evade aging. Aging vol. 7 334–339 https://www.aging-us.com/article/100744/text (2015).

45. Horvath, S. & Levine, A. J. HIV-1 Infection Accelerates Age According to the Epigenetic Clock. J. Infect. Dis. 212, 1563–1573 (2015).

46. Levine, M. E. et al. Menopause accelerates biological aging. Proc. Natl. Acad. Sci. 113, 9327–9332 (2016).

47. Singh, S. S. et al. Association of the IgG N-glycome with the course of kidney function in type 2 diabetes. BMJ Open Diabetes Res. Care 8, e001026 (2020).

48. Lemmers, R. F. H. et al. IgG glycan patterns are associated with type 2 diabetes in independent European populations. Biochim. Biophys. Acta BBA-Gen. Subj. 1861, 2240–2249 (2017).

49. Menni Cristina et al. Glycosylation Profile of Immunoglobulin G Is Cross-Sectionally Associated With Cardiovascular Disease Risk Score and Subclinical Atherosclerosis in Two Independent Cohorts. Circ. Res. 122, 1555–1564 (2018).

50. Belsky, D. W. et al. Eleven Telomere, Epigenetic Clock, and Biomarker-Composite Quantifications of Biological Aging: Do They Measure the Same Thing? Am. J. Epidemiol. 187, 1220–1230 (2017).

51. Liu, Z., Leung, D. & Levine, M. Comparative analysis of epigenetic aging clocks from CpG characteristics to functional associations. 29.

52. Li, X. et al. Longitudinal trajectories, correlations and mortality associations of nine biological ages across 20-years follow-up. eLife 9, e51507 (2020).

53. Lee, Y. et al. Blood-based epigenetic estimators of chronological age in human adults using DNA methylation data from the Illumina MethylationEPIC array. BMC Genomics 21, 747 (2020).

54. Lu, A. T. et al. DNA methylation GrimAge strongly predicts lifespan and healthspan. Aging 11, (2019).

55. McCrory, C. et al. GrimAge outperforms other epigenetic clocks in the prediction of age-related clinical phenotypes and all-cause mortality. J. Gerontol. Ser. A doi:10.1093/gerona/glaa286.

56. Puciċ, M. et al. High throughput isolation and glycosylation analysis of IgG-variability and heritability of the IgG glycome in three isolated human populations. Mol. Cell. Proteomics MCP 10, M111.010090–M111.010090 (2011).

57. Klariċ, L. et al. Glycosylation of immunoglobulin G is regulated by a large network of genes pleiotropic with inflammatory diseases. Sci. Adv. 6, eaax0301 (2020).

58. Leitsalu, L. et al. Cohort Profile: Estonian Biobank of the Estonian Genome Center, University of Tartu. Int. J. Epidemiol. 44, 1137–1147 (2015).

59. Smith, B. H. et al. Cohort Profile: Generation Scotland: Scottish Family Health Study (GS:SFHS). The study, its participants and their potential for genetic research on health and illness. Int. J. Epidemiol. 42, 689–700 (2013).

60. Sudlow, C. et al. UK Biobank: An Open Access Resource for Identifying the Causes of a Wide Range of Complex Diseases of Middle and Old Age. PLOS Med. 12, e1001779 (2015).

61. Min, J. L., Hemani, G., Davey Smith, G., Relton, C. & Suderman, M. Meffil: efficient normalization and analysis of very large DNA methylation datasets. Bioinformatics 34, 3983–3989 (2018).

62. Aryee, M. J. et al. Minfi: a flexible and comprehensive Bioconductor package for the analysis of Infinium DNA methylation microarrays. Bioinformatics 30, 1363–1369 (2014).

63. Yang, J., Lee, S. H., Goddard, M. E. & Visscher, P. M. GCTA: A Tool for Genome-wide Complex Trait Analysis. Am. J. Hum. Genet. 88, 76–82 (2011).

64. Dhingra, R. et al. Evaluating DNA methylation age on the Illumina MethylationEPIC Bead Chip. PLOS ONE 14, e0207834–e0207834 (2019).

65. Sales, S. et al. Gender, Contraceptives and Individual Metabolic Predisposition Shape a Healthy Plasma Lipidome. Sci. Rep. 6, 27710 (2016).

66. Leek, J. T. svaseq: removing batch effects and other unwanted noise from sequencing data. Nucleic Acids Res. 42, 161–161 (2014).

67. Lundberg, M., Eriksson, A., Tran, B., Assarsson, E. & Fredriksson, S. Homogeneous antibody-based proximity extension assays provide sensitive and specific detection of low-abundant proteins in human blood. doi:10.1093/nar/gkr424.

68. Long, T. et al. Whole-genome sequencing identifies common-to-rare variants associated with human blood metabolites. Nat. Genet. 49, 568–578 (2017).

69. Friedman, J., Hastie, T. & Tibshirani, R. Regularization Paths for Generalized Linear Models via Coordinate Descent. J. Stat. Softw. 33, 1–22 (2010).

70. Kim, S. ppcor: An R Package for a Fast Calculation to Semi-partial Correlation Coefficients. Commun. Stat. Appl. Methods 22, 665–674 (2015).

71. Cox, D. R. Regression Models and Life-Tables. J. R. Stat. Soc. Ser. B Methodol. 34, 187–220 (1972).

