## Supplementary Information for "A catalogue of omics biological ageing clocks reveals substantial commonality and associations with disease risk"

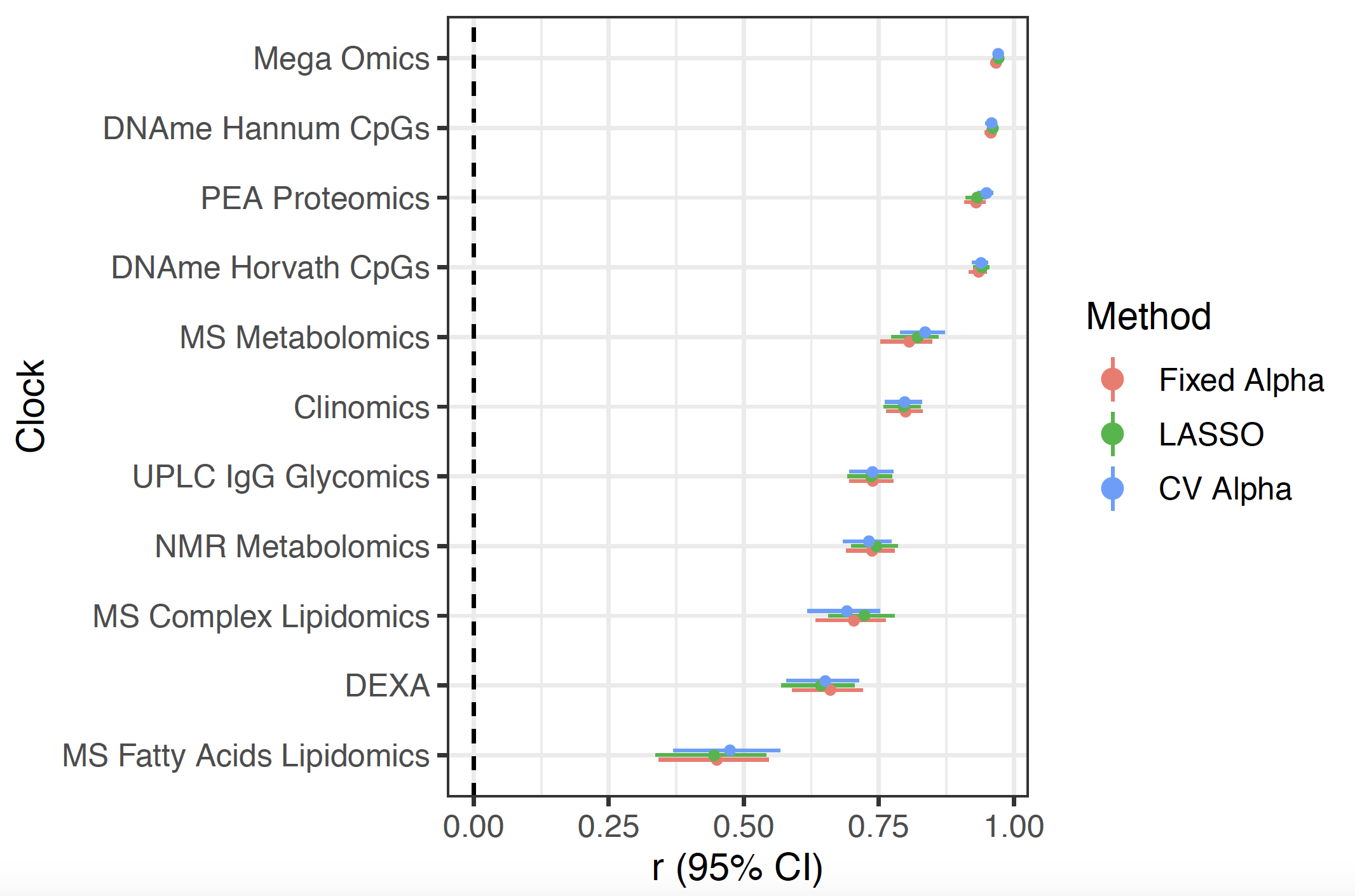

**Supplementary Figure 1. Correlation of ChronAge and OCA were consistent, independent of penalised regression method.** Correlation (r) with 95% of confidence intervals of chronAge with omics clock estimated age (OCA) indicated on the y-axis via elastic net regression with a fixed alpha of 0.5, cross validated alpha and LASSO regression in the ORCADES testing sample.

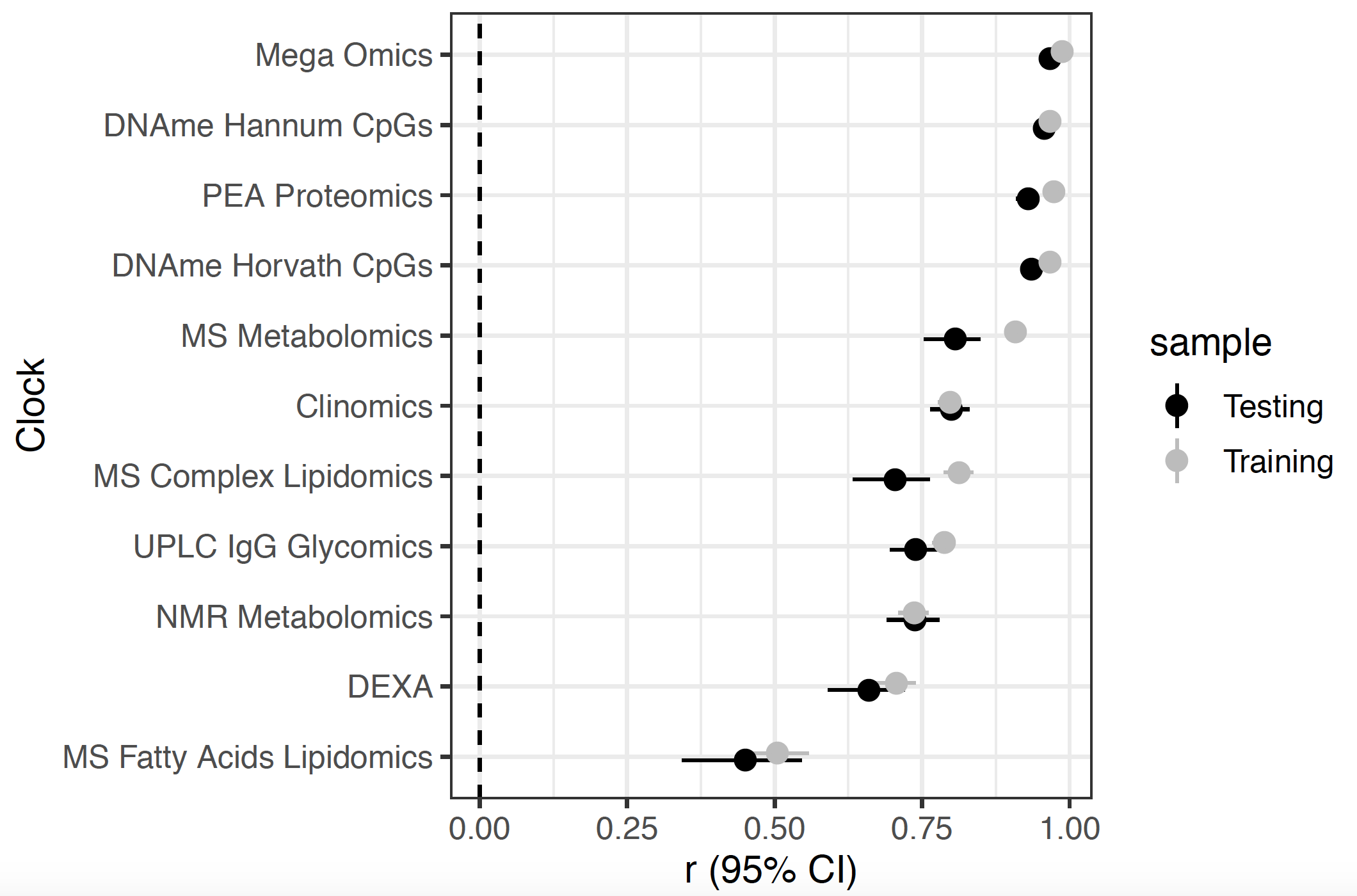

**Supplementary Figure 2. Correlation of ChronAge and OCA in ORCADES Training and Testing Samples.** Correlation (r) with 95% of confidence intervals of chronAge with OCA indicated on the y-axis in the ORCADES Training and Testing samples.

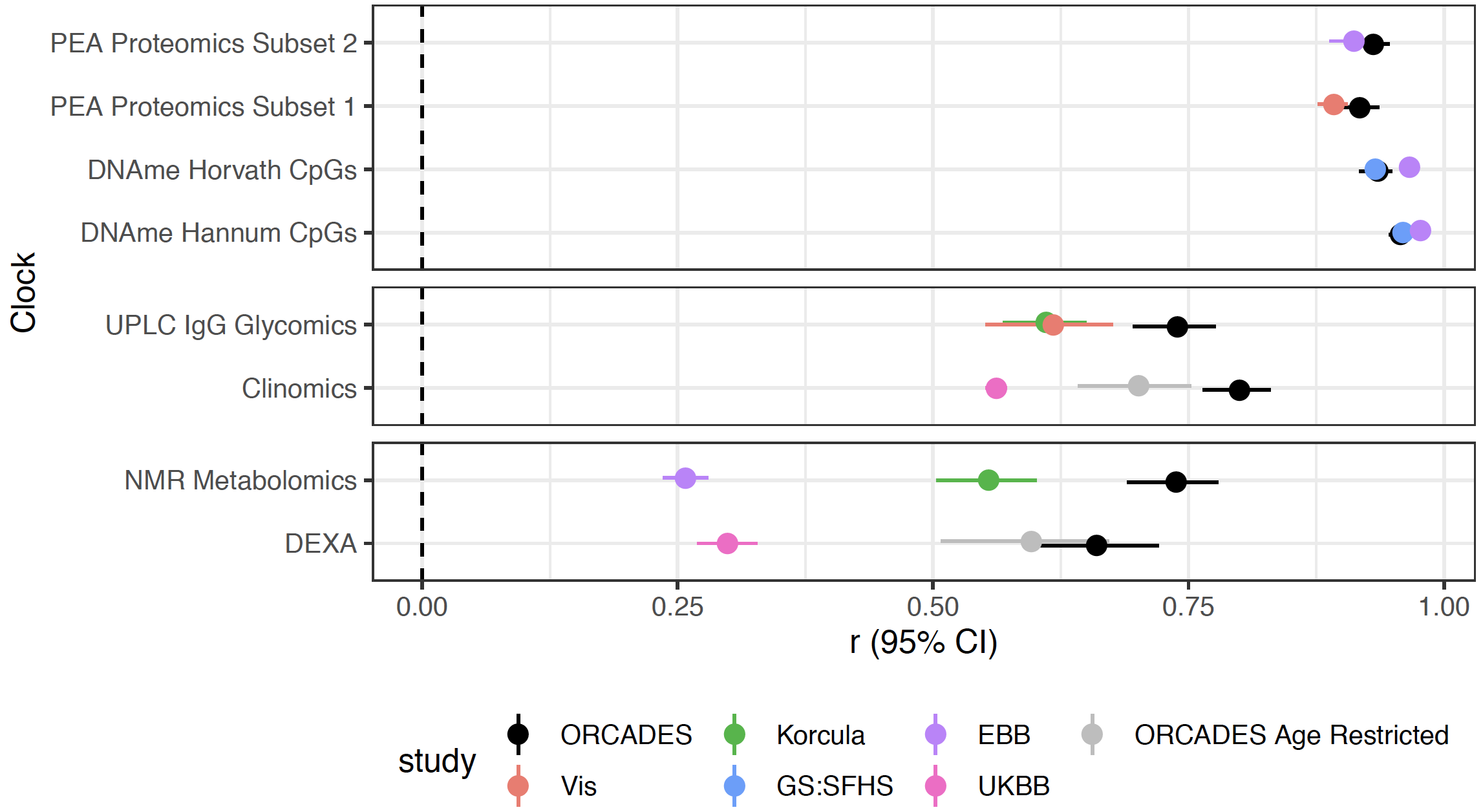

**Supplementary Figure 3. Omics clocks trained in ORCADES predict chronAge in unrelated cohorts.** The correlation of OCA with ChronAge (x-axis) by the specified clock (y-axis). With the correlation in the ORCADES testing sample in black and additional populations as specified. The correlation in a restricted age range (40-75) ORCADES testing sample is shown in comparisons involving the UKBB shown in grey.

We found clocks built using the subsets of PEA proteomics measures available in our validation cohorts correlating with chronAge nearly as highly in Croatia-Vis (r=0.89) and EBB (r=0.91) as in the ORCADES testing sample (r=0.91 and r=0.93). Similarly, both Hannum and Horvath CpG based clocks achieved comparable correlations between OCA and chronAge in EBB (Hannum: r=0.98, Horvath: r=0.97) and GS:SHFS (Hannum: r=0.96, Horvath r=0.93) as in the ORCADES testing sample (Hannum: r=0.96, Horvath r=0.93). The UPLC IgG glycomics and Clinomics OCA were still correlated with chronAge in independent cohorts (UPLC IgG glycomics: r= 0.62 Croatia-Vis, r=0.61 Croatia-Korcula, Clinomics: r=0.56 UKBB) but less than in the ORCADES testing sample (UPLC IgG glycomics: r=0.74, Clinomics: r=0.80). There was correlation between NMR metabolomics estimated and chronAge in Croatia-Korcula, r=0.55 compared to r=0.73 in ORCADES however only a correlation of r=0.26 in EBB. Similarly, we found that the DEXA estimated age in UKBB correlated substantially lower with chronAge than in ORCADES (UKBB: r=0.30, ORCADES: r=0.66).

To assess whether the poor correlation of DEXA OCA and chronAge in UKBB was due to the difference in the ranges of chronAge of individuals in ORCADES compared to the UKBB we also compared a clock that was evaluated in ORCADES individuals between 40-75 (the recruiting age range of UKBB, compared to the 16-100 in the full ORCADES dataset). Despite the DEXA OCA having a lower correlation with chronAge in the age restricted ORCADES sample, r=0.60 compared with r=0.66 in the full age range sample, it is still drastically higher than the r=0.30 found in UKBB.

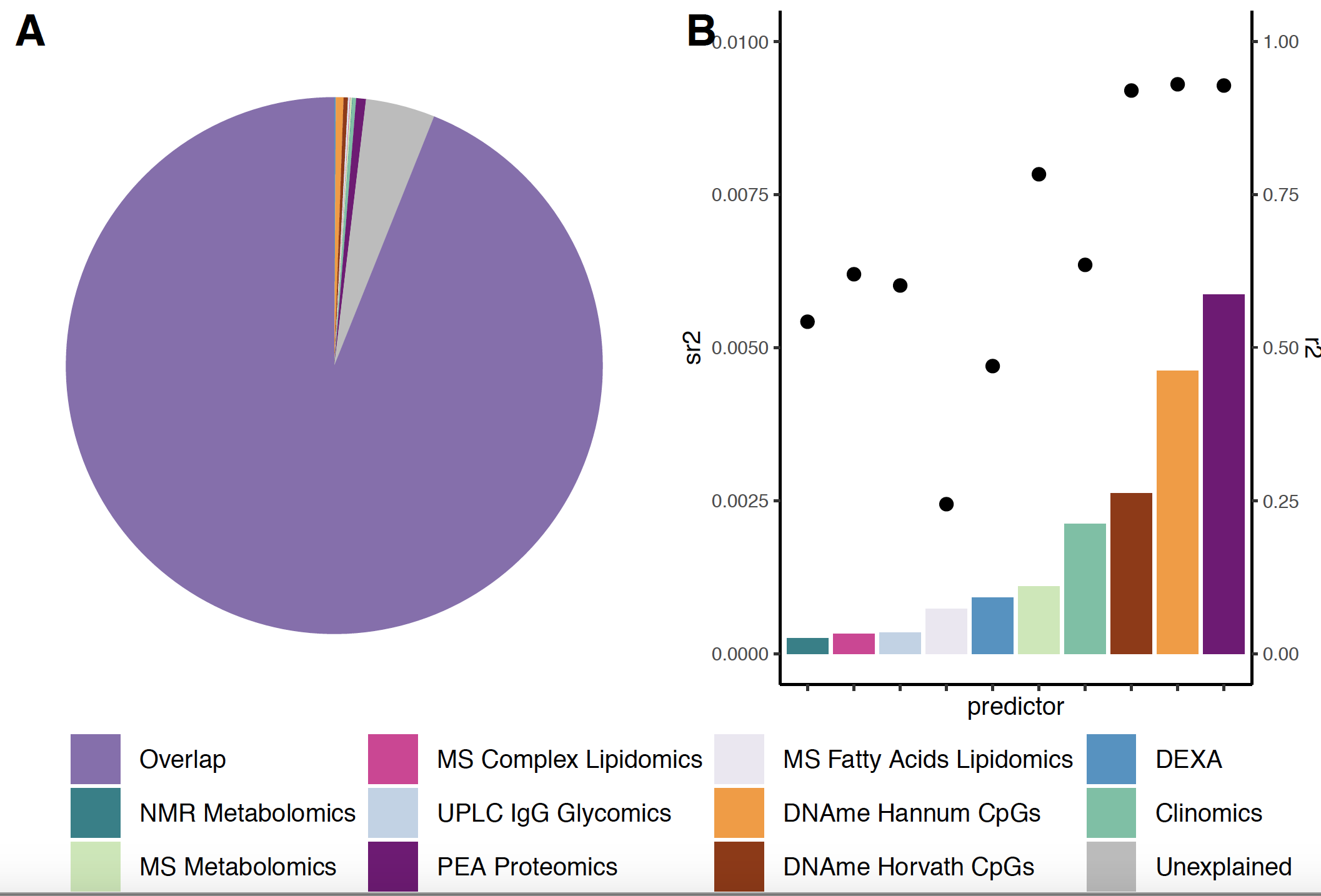

**Supplementary Figure 4. Overlapping and unique variance in ChronAge explained across 10 omics clocks**. A) Partition of variance in ChronAge explained into that explained by 2 or more clocks (overlap), that not explained by any clock (unexplained), and that explained by each of the 10 clocks uniquely. Segments coloured by component explaining the variance in chronAge. B) squared part correlations (sr^2^) (bars): unique variance in chronAge explained by each of the 10 clocks from figure A on the left-hand y-axis. R^2^ (points) indicate the total variance explained in chronAge by each clock (right hand y-axis).

Interestingly, the proportion of unique variance in chronAge explained by each OCA does not entirely mirror the univariate
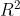
 (black dots). It is important to note that the similarity between assays likely influences the proportion of unique variance in chronAge explained (at its most extreme, were a clock duplicated, it would explain no unique variance). This may explain why NMR metabolomics and MS complex lipidomics clocks have some of the lowest proportions of unique variance explained, despite NMR metabolomics and MS complex lipidomics having an R^2^ higher than DEXA OCA and comparable to Clinomics. Interestingly, the DEXA and MS fatty acids lipidomics OCAs explain more unique variance than several clocks with higher R^2^.

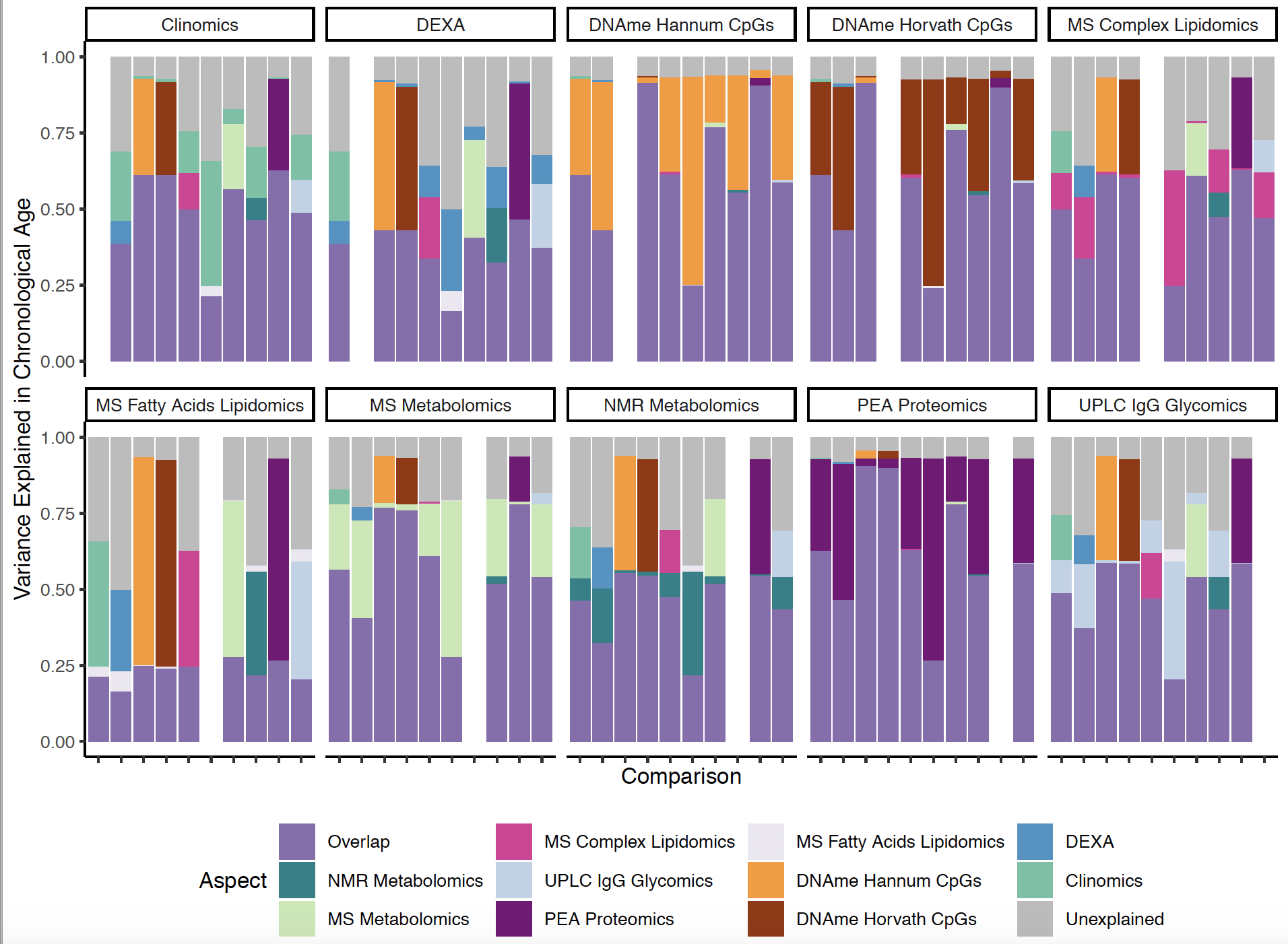

**Supplementary Figure 5. Pairwise Comparisons of Variance Explained in ChronAge**. Pairwise comparison of variance in chronAge explained by OCA of the pairs of clocks in ORCADES. Comparison indicated on the x-axis, with the variance in chronAge explained on the y-axis. The colour of the bar indicates the aspect explaining the variance. For each comparison the proportion of variance explained by both clocks in the comparison (Overlap), the variance that remains unexplained fitting a bivariate model (unexplained) and the unique variance in chronAge explained by each of the two clocks in the comparison.

Partly to consider the effect of two similar clocks affecting the unique variance explained, we performed pairwise comparisons, the unique variance in chronAge explained by each clock in the comparison was again calculated as the squared part correlation while controlling for the other clock in the pair. The overlap indicated is therefore the proportion of variance in chronAge explained by both clocks in the pair. Reiterating the results in Supplementary Figure 4A, Supplementary Figure 5 shows that for 8 out of 10 clocks the mean percentage of variance explained in chronAge by both clocks (the overlap) is greater than 45%. The MS Fatty Acids Lipidomics and DEXA clocks had lower mean overlap, 23.2% and 36.9% respectively. Interestingly clocks that had higher correlations between OCA and chronAge, such as PEA Proteomics and DNAme based clocks were found to be contributing most of the additional variance in chronAge not explained by the overlap of both clocks. Conversely, the MS Fatty Acids Lipidomics clock, the clock with the lowest correlation between OCA and chronAge appears to contribute little of the additional variance in chronAge not already explained by the other clock across all comparisons.

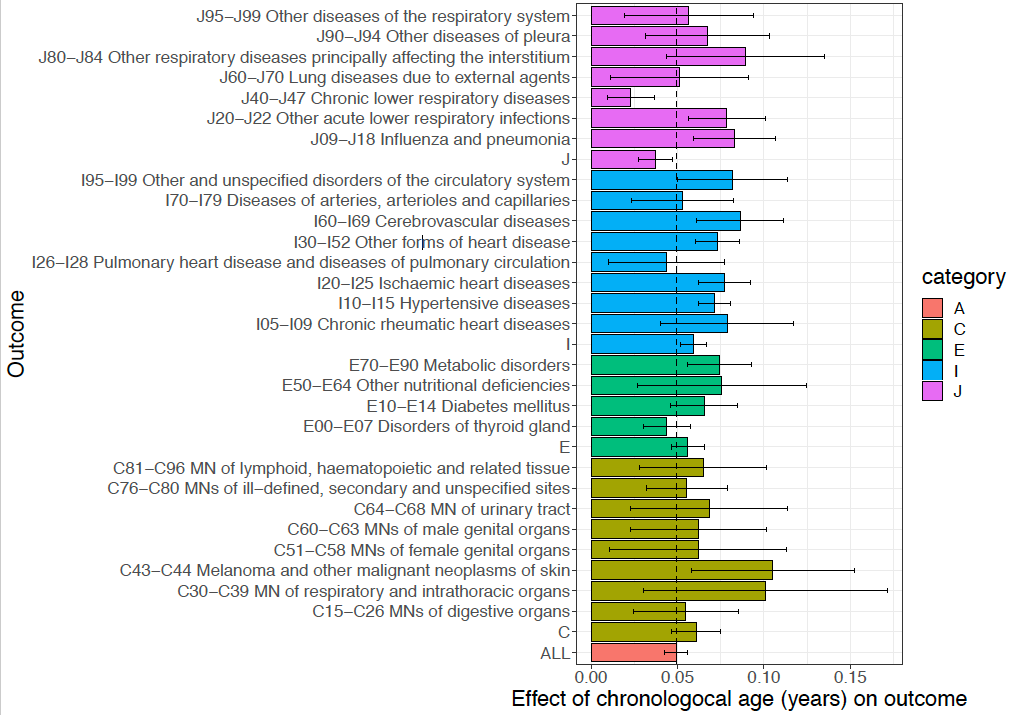

**Supplementary Figure 6a. Associations of disease incidence with chronAge.** Effect and its 95% CI: the log_e_HR of chronAge on the incidence of the disease since participation, using a Cox Model. ICD 10 Chapters (i.e. whole Categories) count the first occurrence (post assessment) of any disease within the letter/category/chapter (including those blocks dropped from the individual block analysis due to lack of power) as incidence. Participants prevalent at assessment (i.e. a recorded prior incidence) within any grouping at assessment were excluded from the analysis of that grouping. The dashed line represents the hazard of age on any occurrence of the disease chapters under consideration, a hazard ratio of 0.0492, representing a doubling of incidence rate every 14 years. Distinctions in observed individual effects sizes from this were (visually) judged more materially due to sampling variance than true effects, and so that single factor was chosen as our best estimate of the age effect on each disease. MNs: Malignant neoplasms Associations are only shown for those disease groups that passed QC and were taken forward to association testing with OCAA.

8/8 risk factors and 43/44 disease groupings associated with chronAge in the expected positive direction, except for cortisol and FEV1 which decline with chronAge. The disease exception, J00-J06 Acute respiratory infections, was not nominally significantly different from zero (log_e_HR/SE -0.025/0.017). All the risk factors, and 34 of the disease groupings associations were significant after allowing for multiple testing (passed FDR 10% risk factors and diseases considered separately, one sided test H_1_:b>0). For these 34 groupings there thus was reasonable power to detect associations with chronAge and so potentially biological OCAA. 2 disease groups had fewer than 5 cases and were excluded from the subsequent analysis, to further limit the burden of multiple testing.

The effect (log_e_HR/SE) of one year of chronAge at outset on the first incidence of any of the diseases was 0.0492/0.00323, a doubling roughly every 14 years. This pattern was generally similar to the estimated effects for each disease individually, noting these are on the same (logistic) scale. With the largest observed differences arising from diseases with larger standard errors. However, the effect (log_e_HR/SE) of one year of chronAge on the risk factors varied more, although again they were on the same (standardised) scale. FRS and FEV1 (0.054/0.00083 and -0.041/0.00088) were most sensitive, whilst CRP and creatinine were less sensitive (effect/SE of 1 year of chronAge on standardised trait 0.0092/0.0012 and 0.0090/0.0015 respectively) as shown in Supplementary Figure 6b, whilst standard errors of the effect sizes were generally smaller (as a proportion of the effect).

The remaining 32 disease groups along with all the risk factors were taken forward to association testing with OCAA, using the same models, with age and sex as covariates. Power was expected to be lower, due to lower variation and attenuation in OCAA compared to chronAge. As our principal purpose was to examine the effect of OCAA compared to chronAge, effect sizes of OCAA were rescaled so that the effect of one year of chronAge was one. This was done by dividing the observed effect of OCAA by the effect of chronAge on the outcomes. Given the similarities of the chronAge effects (and wide SEs) for diseases, this was done using the single factor 0.049187 log_e_HR. Whereas for risk factors this was done trait-by-trait (the effect of chronAge on the single trait) as these effects varied more and had lower standard errors.

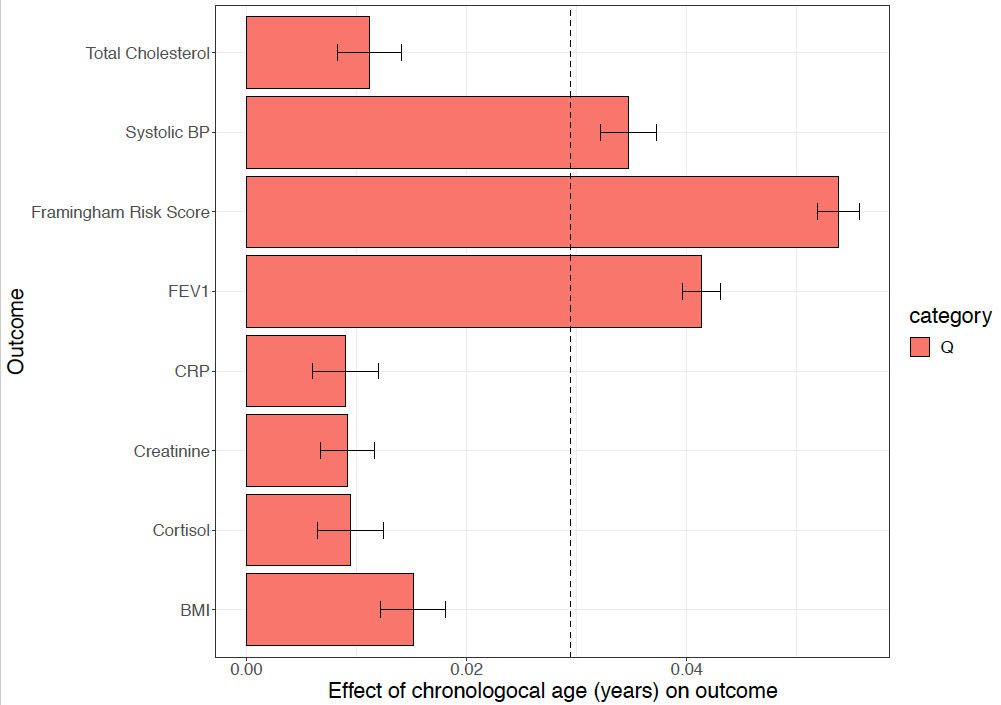

**Supplementary Figure 6b. The strength of associations of risk factors with chronAge varies.** FEV1: Forced expiratory volume one second, CRP: C-reactive Protein, BMI: Body Mass Index. Effect: the estimated increase (and 95% CI) in standardised trait per year of chronAge using a linear model, with sex as a covariate. Traits which decrease as age increases (FEV1, cortisol) have been converted to ageing traits, by reversing their signs.

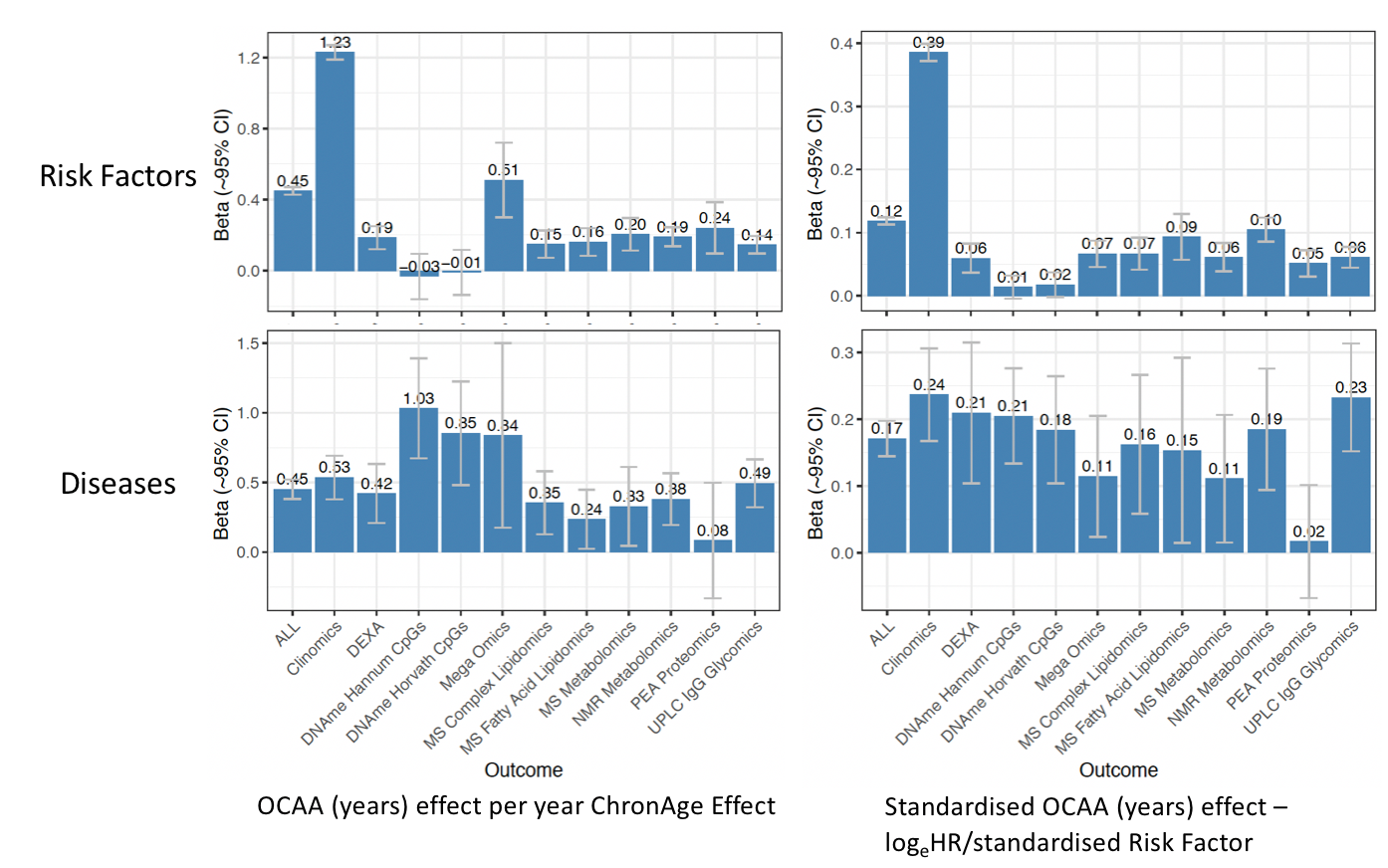

**Supplementary Figure 7a. Averaged effects of OCAA across diseases and risk factors.**

The Left-hand side shows the effect of OCAA in years per year of chronAge effect (OCAA effect divided by chronAge effect) IVW averaged across outcomes (either risk factors or diseases as specified on the y-axis). The right-hand side shows the effect of standardised OCAA (units of phenotypic standard deviation) IVW averaged across outcomes.

Beta: the observed effect of OCAA on outcome. Beta was IVW averaged across outcomes.

SEs were calculated as the inverse root sum of the precisions (not strictly valid given correlated tests). Error bars shown are +/- 2SEs. OCAA: omics clock estimated age acceleration

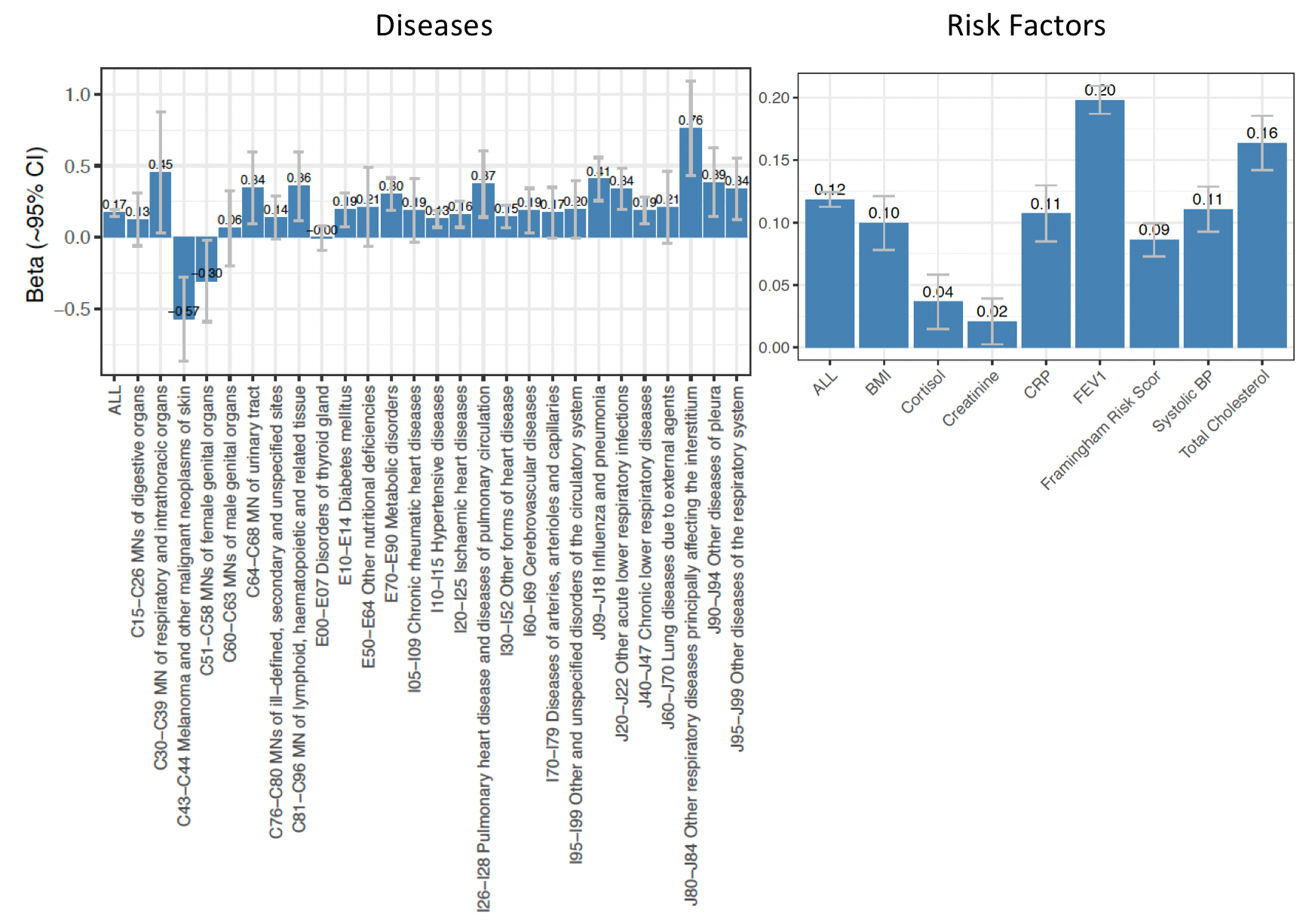

**Supplementary Figure 7b. Average effect across clocks of standardised OCAA upon outcome.** Beta: the observed effect of OCAAs on outcome. Beta was IVW averaged across OCAAs. SEs were calculated as the inverse root sum of the precisions (not strictly valid given correlated tests). Error bars shown are +/- 2SEs. OCAA: omics clock estimated age acceleration

[a] Diseases

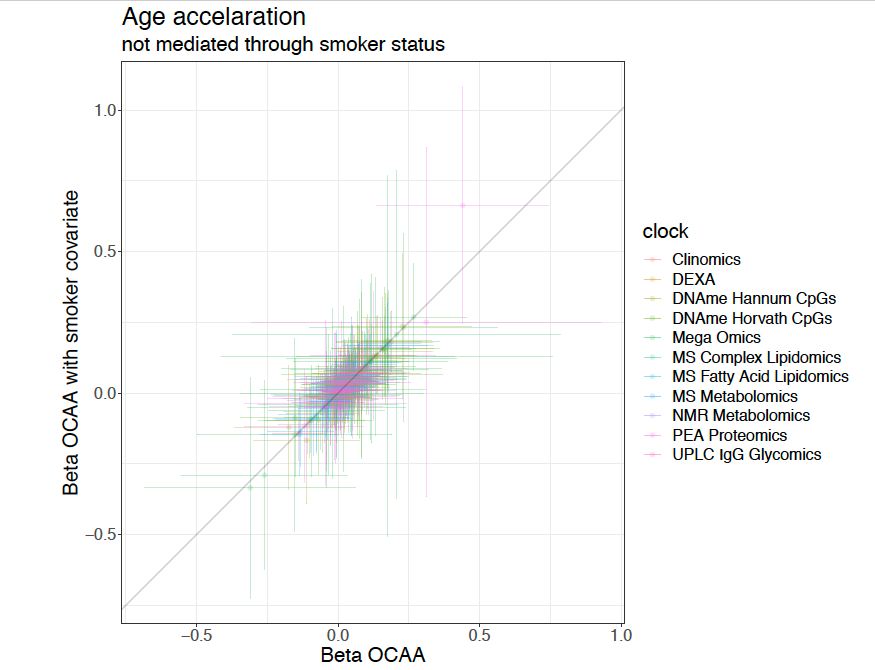

[b Risk Factors]

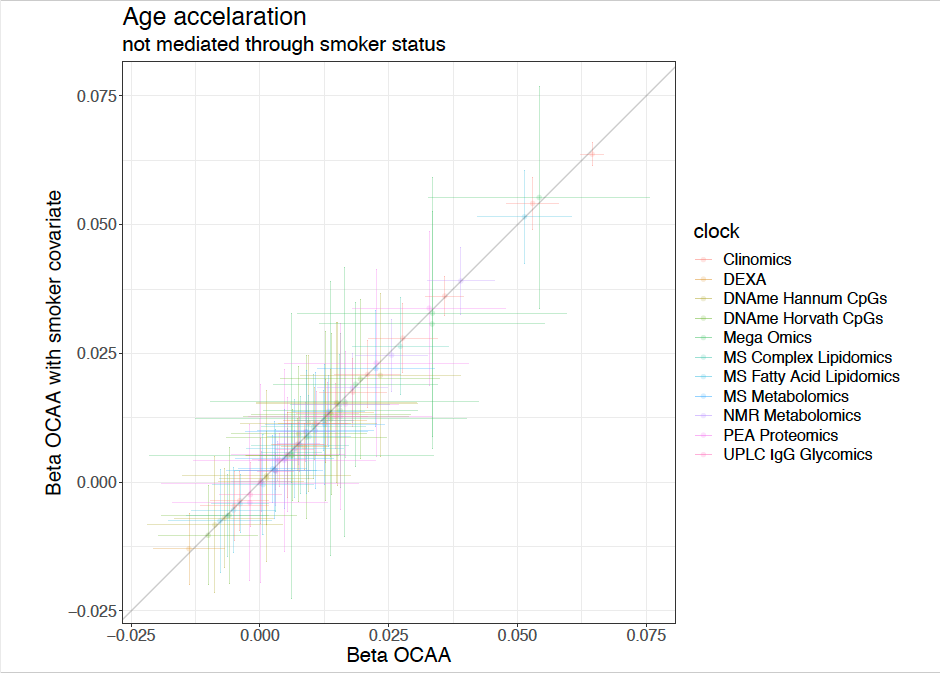

**Supplementary Figure 8. Fitting smoking as a covariate does not appear to materially affect the association between OCAA and outcomes.**

Beta OCAA - the observed effect of OCAA on the outcome under the models (see main text). Beta OCAA with smoker covariate - the observed effect of OCAA on the outcome under the same model, but with smoking fitted.

Across all the associations studied for 11 clocks against 32 diseases and 8 risk factors, we found that the IVW ratio of the estimated effect of OCAA with and without smoking fitted as a covariate were 1.023 and 1.008 respectively. Individual test p-values for the ratio of the effects not being one all exceeded 0.4. Visual analysis confirmed these results: that smoking was not a material confounder of health-OCAA associations.

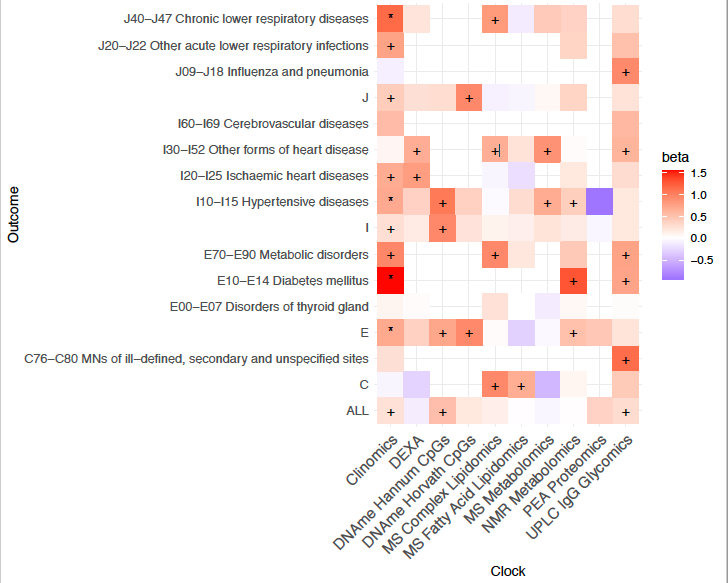

**Supplementary Figure 9a. OCAA positively (pink), and often significantly, associates with disease incidence in most cases where there is reasonable power**. +/* Association nominally(p<5%)/FDR10% significant in the frequentist test of H1:b>0 (FDR is determined across all tests shown in STXX, not just those shown here). Beta: the relative effect of a year of OCAA to a year on chronAge on outcome (measured in log_e_ hazard ratios). A value of one means the estimate of the effect of chronAge and OCAA are the same. Clock: the omics clock on which OCAA was measured. The mega-omics clock is not shown as it never met the SE<0.5 criterion. Disease group: the set of diseases (defined by ICD 10 codes) which were tested for first incidence after assessment against the clock (already prevalent cases were excluded).

The presence of an entry in this figure denotes power, whilst its intensity denotes the size of the effect. Clinomics OCAA is thus relatively powerful and has large effects, as does the UPLC IgG Glycomics OCAA, albeit to a lesser extent. The more accurate clocks at estimating chronAge such as DNAme and PEA Proteomics based clocks on the other hand, show less power, although the Horvath CpGs clock does show some reasonably strong effect sizes.

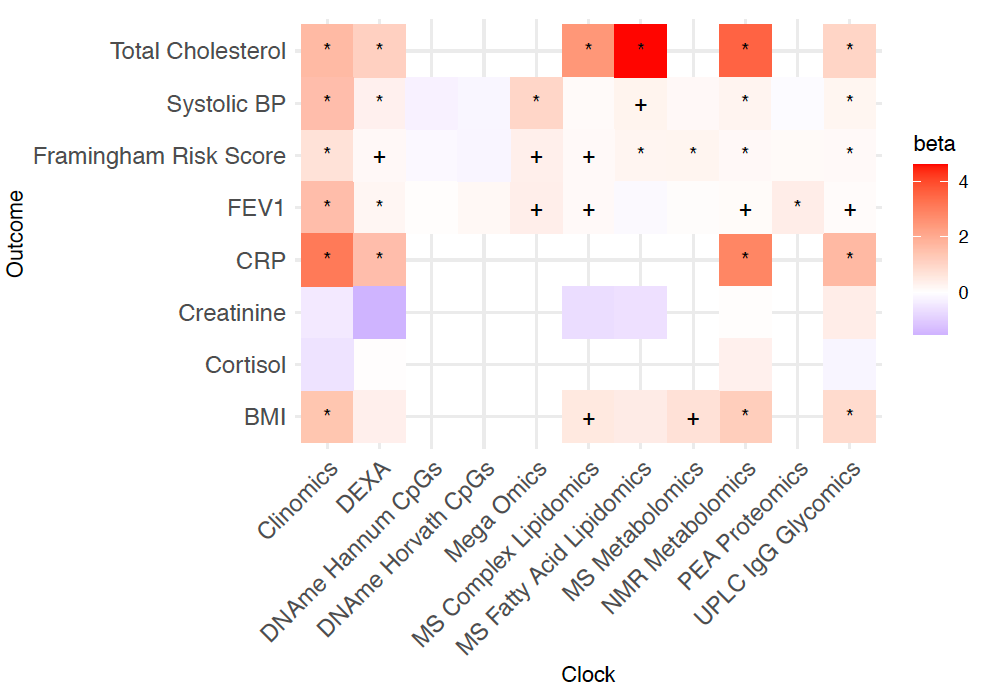

**Supplementary Figure 9b. OCAA positively (pink) and often significantly associates with risk factors in most cases where there is reasonable power**. +/* Association nominally(p<5%)/FDR10% significant in the frequentist test of H1:b>0 (FDR is determined across all tests shown in STXX_q, not just those shown here). Beta: the relative effect of a year of OCAA to a year on chronAge on outcome. A value of one means the estimate of the effect of chronAge and OCAA are the same.

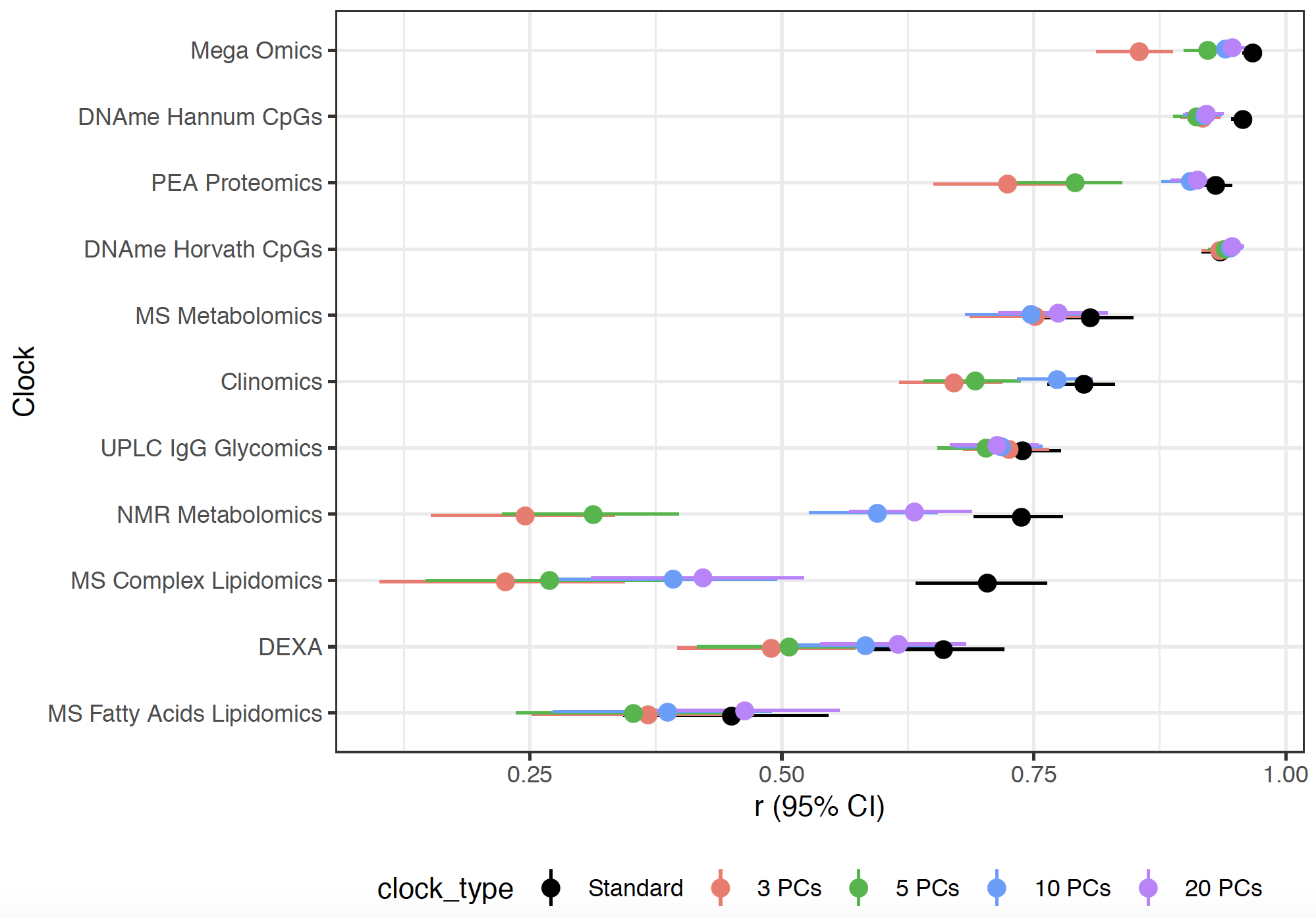

**Supplementary Figure 10. Correlation of chronAge and OCA from clocks built using 3, 5, 10, 20 PCs**. Correlation (r) and 95% confidence interval of chronAge and OCA indicated on the y-axis using models constructed from 3, 5, 10 and 20 principal components of the assay in the ORCADES testing sample compared to the standard clock (black).

[a - Risk Factors]

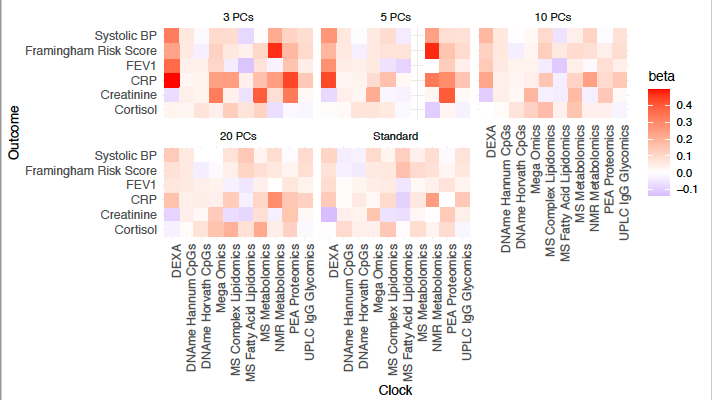

**Supplementary Figure 11a. Reducing dimensionality of omics dataset used to build clocks increases the predictive ability of OCAA for risk factors**

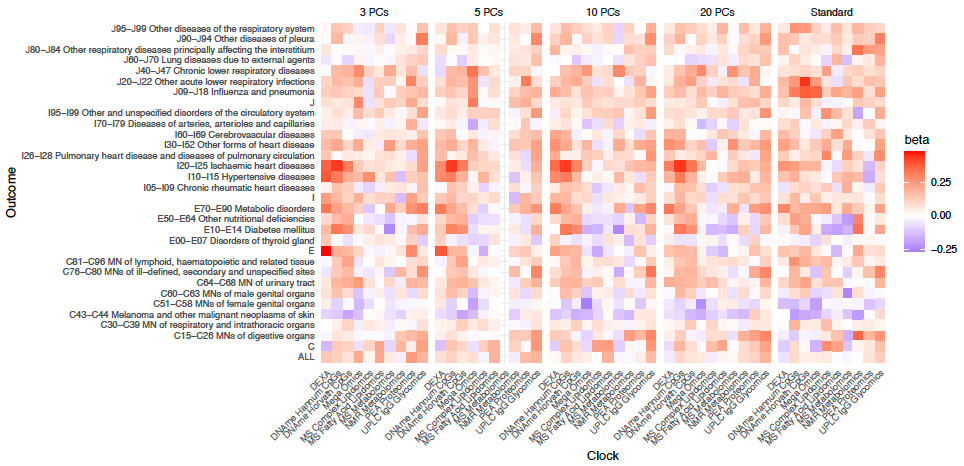

**Supplementary Figure 11b. Reducing dimensionality of omics dataset to train ChronAge makes little difference to the predictive ability of OCAA for risk factors.** Beta: the effect of a year of standardised (within clock) OCAA on outcome (measured in log_e_ hazard ratios, for diseases, effect sizes for standardised risk factors). Estimates were shrunk using a prior to reduce the possibility that frequentist best estimate beta was predominantly a consequence of a large SE. Clock: the omics clock on which OCAA was measured. Disease group: the set of diseases (defined by ICD 10 codes) which were tested for first incidence after assessment against the clock (already prevalent cases were excluded). Cholesterol/BMI which showed a particularly large effect from MS Fatty Acids Lipidomics/DEXA OCAA, excluded here to aid visualisation. X PCs: the number of PCs of the omic used as predictors to create the chronAge and OCAA measures.

OCAAs (and risk factors) were standardised, whilst hazards were left on their log_e_ scale. The resultant measures of the effect of OCAA on outcome gave a measure of the ability of the OCAA to distinguish amongst individuals.

Clinomics was excluded from this analysis as it was based on only 12 predictors. We continued to exclude Total Cholesterol, but also excluded BMI as too close in nature to some of the predictors used.
